# Lumican regulates cervical corticospinal axon collateralization via non-autonomous crosstalk between distinct corticospinal neuron subpopulations

**DOI:** 10.1101/2021.03.26.437104

**Authors:** Yasuhiro Itoh, Vibhu Sahni, Sara J. Shnider, Jeffrey D. Macklis

## Abstract

Corticospinal neurons (CSN) are the cortical projection neurons that innervate the spinal cord and some brainstem targets with segmental precision to control voluntary movement of specific functional motor groups, limb sections, or individual digits. CSN subpopulations exhibit striking axon targeting specificity from development into maturity: Evolutionarily newer rostrolateral CSN exclusively innervate <u>b</u>ulbar-<u>c</u>ervical targets (CSN_BC-lat_), while evolutionarily older caudomedial CSN (CSN_medial_) are more heterogeneous, with distinct subpopulations extending axons to either bulbar-cervical or thoraco-lumbar segments. However, molecular regulation over specificity of CSN segmental target innervation is essentially unknown. The cervical cord, with its evolutionarily enhanced precision of forelimb movement, is innervated by multiple CSN subpopulations, suggesting inter-neuronal interactions in establishing cervical corticospinal circuitry. Here, we identify that Lumican, previously unrecognized in axon development, controls the balance of innervation between CSN_BC-lat_ and CSN_medial_ within the cervical spinal cord. Remarkably, Lumican, an extracellular matrix protein expressed by CSN_BC-lat_, non-cell-autonomously suppresses axon collateralization in the cervical cord by CSN_medial_. Intersectional viral labeling and mouse genetics further identify that Lumican controls axon collateralization by multiple CSN subpopulations in caudomedial sensorimotor cortex. These results identify inter-axonal molecular crosstalk between CSN subpopulations as a novel mechanism controlling corticospinal circuitry, target density, and competitive specificity. Further, this mechanism has potential implications for evolutionary diversification of corticospinal circuitry with finer scale precision.

## Introduction

Corticospinal neurons (CSN, and related cortico-brainstem neurons; together “CSN”) extend axon projections to subcerebral targets, and make synaptic connections with circuits in the brainstem and spinal cord (Kalil and Dent, 2014; Levine et al., 2012; Sahni et al., 2020). CSN axons thus form the corticospinal tract (CST), the major motor output pathway from the cerebral cortex, essential for voluntary motor control, and the principal circuit underlying skilled movement (Lemon, 2008; Martin, 2005). The vast majority of CSN ultimately control voluntary movements by coordinately moving target muscles of specific functional motor groups, limb sections, or individual digits (Levine *et al*., 2012). The repertoire of skilled movement, such as speech and precise forelimb/digit movement, has expanded significantly through mammalian evolution, and this dramatic expansion is associated with a concomitant evolutionary increase in cortical projections to circuits in the brainstem and spinal cord. This includes both increased numbers of corticospinal axons innervating these subcerebral targets, and expansion of the cortical territory that gives rise to these projections (Dum and Strick, 2002; Lemon, 2008; Schieber, 2007). The ultimate execution of such precise fine motor control necessitates that subcortical circuits receive diverse, yet precise inputs from distinct cortical areas (Dum and Strick, 2002; Lemon, 2008; Schieber, 2007). How these cortical efferents establish diverse yet precise innervation of segmentally-specific spinal and brainstem circuitry during development remains essentially unknown.

Mechanisms controlling broad CSN development and axon targeting have begun to be identified and systematized. Molecular controls have been elucidated that govern the specification and differentiation of broad, hodologically distinct subtypes of neocortical projection neurons— CSN/subcerebral projection neurons (SCPN; CSN are a subset of SCPN), corticothalamic projection neurons (CThPN), and intracortical callosal projection neurons (CPN) (reviewed in (Greig et al., 2013; Ozkan et al., 2020)). CSN differentiation reflects a coordination of developmental programs that operate along temporal, subtype, and areal axes (Greig *et al*., 2013), and these molecular controls together specify and differentiate CSN from other hodologically distinct projection neuron subtypes within the neocortex (Greig *et al*., 2013; Ozkan *et al*., 2020). CSN differentiation is also coordinated via interplay between CSN and other cell types (Thion et al., 2018; Ueno et al., 2013). However, molecular regulation over how CSN developmentally acquire precise segmental target specificity within the rostro-caudal extent of the brainstem and spinal cord remains essentially unknown.

Importantly, CSN exhibit striking anatomical and functional diversity (Donoghue and Wise, 1982; Liu et al., 2018; Neafsey et al., 1986; Penfield and Rasmussen, 1950; Schieber, 2007; Steward et al., 2020; Tennant et al., 2011; Ueno et al., 2018; Wang et al., 2017; Woolsey et al., 1952)— some CSN extend axons to targets in the brainstem and cervical spinal cord to control face and arm movement, while others extend axons to thoraco-lumbar segments to control trunk and leg movement. Further, CSN have subsets in multiple cortical areas, spanning far beyond primary motor cortex (M1). CSN in rostrolateral sensorimotor cortex are evolutionarily newer, and exclusively innervate bulbar-cervical targets (referred to here as CSN_BC-lat_). In contrast, CSN in caudomedial sensorimotor cortex, including M1, are evolutionarily older, and are relatively more heterogeneous (referred to here as CSN_medial_), with interdigitated subpopulations extending axons to either bulbar-cervical or thoraco-lumbar spinal cord, maintained from early development into maturity (Kaas, 2004; 2013; Krubitzer, 2007; Sahni et al., 2018). Thus, CSN_medial_ can be further subdivided into CSN_BC-med_ and CSN_TL_, respectively (Sahni *et al*., 2018). We have investigated CSN diversity, and identified that anatomically distinct CSN subpopulations are also molecularly distinct, revealing intrinsic molecular controls over CSN axon extension in the CST to distinct spinal segmental levels (Sahni *et al*., 2018). However, it remains unclear how subsequent axon collateralization by these distinct CSN subpopulations is molecularly regulated (Canty and Murphy, 2008; Chédotal, 2019). Of note, the cervical cord, which coordinates skilled forelimb/arm movements, receives inputs from multiple cortical areas spanning far beyond primary motor cortex (Dum and Strick, 2002; Steward *et al*., 2020; Ueno *et al*., 2018) and is indeed innervated to greater or lesser extent by all three CSN subpopulations mentioned above (CSN_BC-lat_, CSN_BC-med_, and CSN_TL_) (Sahni *et al*., 2018). This raises a fundamental question regarding how distinct CSN subpopulations elaborate the appropriate contribution of axon collaterals into segments of the cervical spinal gray matter, and whether there is a competitive crosstalk between axons of these distinct CSN subpopulations. Further, corticospinal circuitry is known to undergo rewiring in response to injury (Ghosh et al., 2010; Kaiser et al., 2019; Oudega and Perez, 2012). Identifying developmental regulators of corticospinal axon collateralization might inform approaches aimed at rewiring damaged corticospinal circuitry with greater efficiency and precision (Bareyre et al., 2002; Oudega and Perez, 2012).

Lumican belongs to the family of extracellular matrix (ECM) proteins called small leucine-rich proteoglycans (SLRPs). Although Lumican has been studied in a variety of tissues and organs–notably the cornea and connective tissues–in the context of development, immunity, or cancer (Brézillon et al., 2013; Chen and Birk, 2013), its expression or function during mammalian neural development remains relatively unstudied and thus poorly understood (Long et al., 2018).

Lumican emerged from our prior work as a particularly interesting candidate for potentially controlling CSN axon collateralization for several reasons. *Lumican* encodes a twelve leucine-rich repeat (LRR)-containing ECM protein that is conserved across vertebrates (Brézillon *et al*., 2013; Caterson and Melrose, 2018; Chen and Birk, 2013), and many similar ECM proteins have been shown to regulate axon guidance and targeting (Lowery and Van Vactor, 2009; Tessier-Lavigne and Goodman, 1996). Our previous work identified that *Lumican* is expressed by a select subset of CSN, CSN_BC-lat_, during postnatal development, with peak expression in the second postnatal week, when CSN axons are actively collateralizing in the spinal cord (Arlotta et al., 2005; Sahni *et al*., 2018). Further, a previous study identified a unique, nonsynonymous *Lumican* variant in a patient cohort of amyotrophic lateral sclerosis (ALS) (Daoud et al., 2011), a disease characterized by progressive degeneration of CSN and spinal motor neurons (Ravits et al., 2013; Taylor et al., 2016).

In the present investigation, we identify that Lumican is expressed specifically by CSN_BC-lat_ in the developing mouse cortex, and controls axon collateralization by CSN_medial_ (both CSN_BC-med_ and CSN_TL_) in the cervical cord in a non-cell-autonomous, crosstalk manner. Our data identify a new form of axon guidance mechanism controlling arborization of CSN axon collaterals at specific segmental levels of the spinal cord. This mechanism additionally highlights a novel form of inter-subpopulation control over the establishment of balanced corticospinal innervation, whereby a secreted molecule generated by one CSN subpopulation mediates inter-axonal expulsion and competitive specificity between CSN subpopulations that are thought to be evolutionarily distinguished, potentially generalizable to other subtypes of neurons and other circuitry. Such mechanisms likely represent a key step for establishing precision of corticospinal circuitry, an axon projection pathway in which distinct subpopulations likely controlling distinct functional outputs reside in an interdigitated and non-topographic manner. Our results and their implications might also elucidate how evolutionarily newer CSN integrate into evolutionarily pre-existing circuitry composed of evolutionarily older CSN, and how this enables diversification of circuitry for potential execution of more-refined function.

## Results

### Lumican is specifically and transiently expressed by CSN_BC-lat_ in postnatal mouse cortex

We originally identified that *Lumican* is specifically expressed postnatally by CSN when compared with CPN (Arlotta *et al*., 2005). In more recent work (Sahni *et al*., 2018), we used transcriptional profiling to identify genes differentially expressed between developing CSN_BC-lat_ and CSN_medial_, purified respectively from rostrolateral versus caudomedial sensorimotor cortex in mice (Fig. 1A). In this work, we identified *Lumican* as strikingly differentially expressed by CSN_BC-lat_ compared with CSN_medial_ (Fig. 1B), with almost no detectable expression by CSN_medial_. *Lumican* differential expression by CSN_BC-lat_ steadily increases from postnatal day (P) 1 to P7 (Fig. 1B). These results suggested the hypothesis that Lumican might function in development and/or specificity of circuitry between cortex and bulbar-cervical targets. In particular, Lumican’s known molecular family, characteristics as a secreted proteoglycan, and functions as an ECM protein in other systems suggested its potential roles in regulation of axon growth, and potentially even regulating neurons other than CSN_BC-lat_.

**Figure 1:**
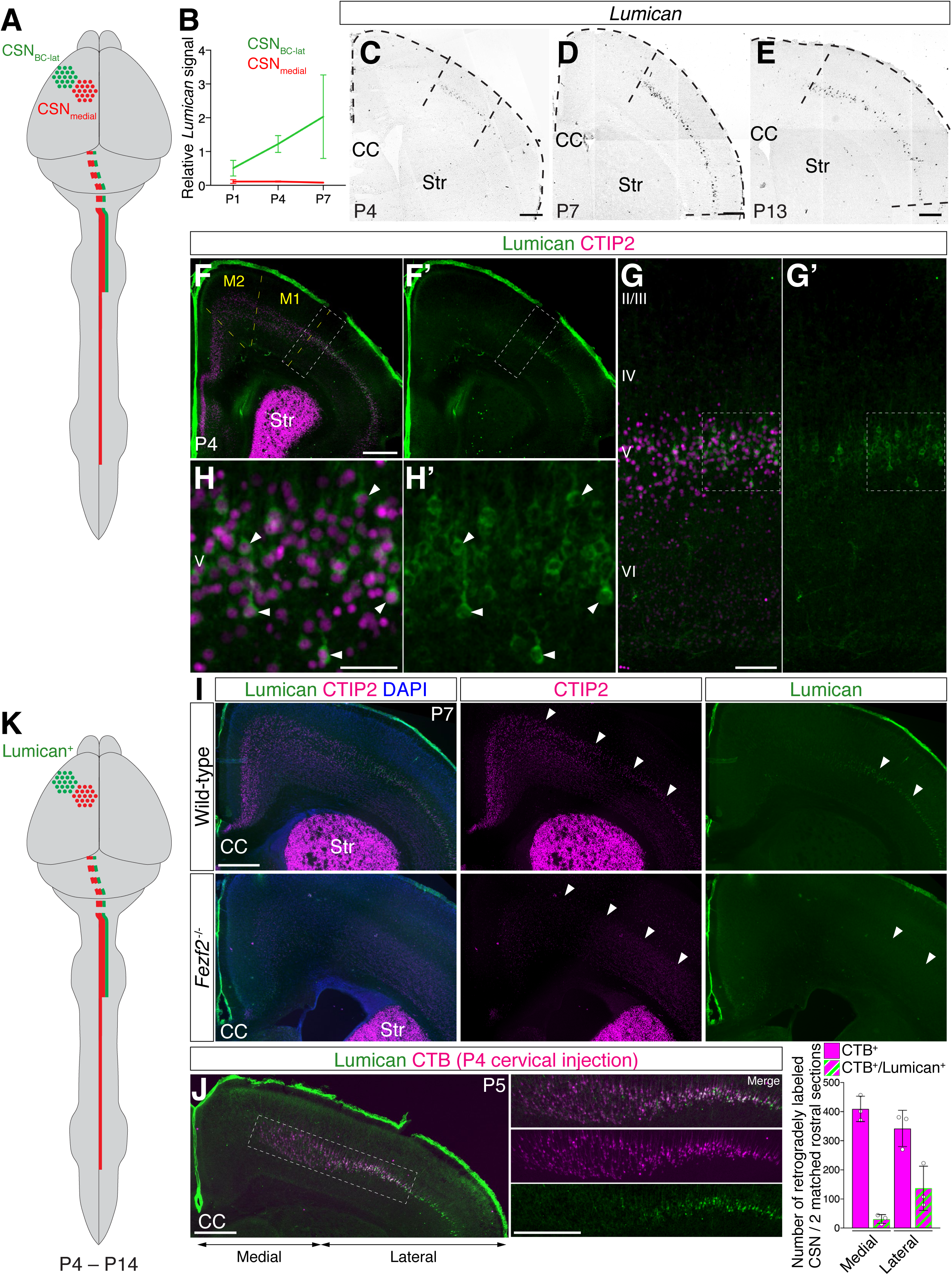
Lumican is expressed by CSN_BC-lat_ in postnatal developing cortex in mice. (A) Schematized representation of developmentally distinct CSN subpopulations. CSN in rostrolateral sensorimotor cortex (CSN_BC-lat_; green) send projections exclusively to brainstem and cervical cord. CSN in caudomedial sensorimotor cortex (CSN_medial_; red) comprise interspersed subpopulations of both bulbar-cervical–projecting CSN and thoraco-lumbar–projecting CSN (Sahni *et al*., 2018). For simplicity, axon collaterals are not illustrated, and CSN are illustrated in only one hemisphere. (B) Temporal profile of *Lumican* expression from microarray analysis at postnatal ages P1, P4, and P7. CSN_BC-lat_, green; CSN_medial_, red. Y-axis represents normalized expression level; data are presented as mean ± SD, n = 2–3. (C–E) *In situ* hybridization on coronal brain sections reveals that *Lumican* is expressed exclusively in lateral cortex at P4, P7, and P13. Scale bars, 400 µm. (F–H) Immunocytochemistry on coronal P4 brain section detects Lumican expression (green) only in lateral cortex, while it is not detected medially (F, F’). Lumican is expressed specifically in layer V (G, G’), and virtually all Lumican-positive cells co-express CTIP2 (magenta) (H, H’). Arrowheads indicate neurons co-expressing Lumican and CTIP2. Scale bars, 500 µm (F); 100 µm (G); 50 µm (H). (I) There is no Lumican expression (green) in P7 *Fezf2* null cortex, which completely lacks CSN and other SCPN, which are labeled by CTIP2 (magenta), while meningeal Lumican expression is preserved. DAPI, blue. Scale bar, 500 µm. (J) Lumican (green) immunocytochemistry on coronal section of P5 brain retrogradely labeled by CTB-555 (magenta) injection into cervical dorsal funiculus at P4. Co-labeling by Lumican and CTB is observed predominantly in lateral cortex, demonstrating Lumican expression selectively by CSN_BC-lat_. Data are presented as mean ± SD, n = 3. Scale bars = 500 µm. (K) Schematized representation of Lumican expression by developing CSN_BC-lat_ (green) between P4 and P14. CC, corpus callosum; M1, primary motor cortex; M2, secondary motor cortex; Str, striatum.

We investigated the temporal course and cell-type-specific expression of Lumican in postnatal neocortex. We first performed *in situ* hybridization at P4, P7, and P13. *Lumican* expression is strikingly restricted to layer V in the lateral sensorimotor cortex, where CSN_BC-lat_ reside, and is excluded from medial sensorimotor cortex, where CSN_medial_ reside. *Lumican* expression in lateral cortex peaks at P7 (Fig. 1C–E). This expression is consistent with the data from the differential gene expression analyses described above, and validates that *Lumican* is expressed by CSN_BC-lat_, but not by CSN_medial_ (Fig. 1B). Immunocytochemical analyses additionally confirm that Lumican expression is confined to layer V in lateral cortex at P4 and P8 (Fig. 1F, G and S1A, B). Importantly, Lumican protein is more abundant rostrolaterally, and absent medially in M1 and secondary motor cortex (M2) areas (Fig. 1F and S1A–C). Neocortical Lumican expression remains restricted to lateral layer V at P14, although at a lower level, and becomes undetectable by P21 (Fig. S1D).

We next investigated whether Lumican is expressed only by SCPN in lateral cortex. The overwhelming majority of Lumican-positive cells (88.1 ± 2.0%) also express high levels of CTIP2, an SCPN-specific control at high expression level (Arlotta *et al*., 2005). A small fraction of Lumican-expressing cells (24.2 ± 5.1%) also express SATB2, which was initially identified as a CPN developmental control (Alcamo et al., 2008; Britanova et al., 2008), but later found to be co-expressed with CTIP2 by a substantial fraction (20–40%) of CTIP2-expressing SCPN earlier in development until co-expression resolves by P15 (Leone et al., 2015) (Fig. 1F–H and S1A). Importantly, the vast majority (95%) of SATB2+/Lumican+ double-positive neurons also express CTIP2. Together, these results strongly indicate that Lumican is expressed specifically by SCPN in rostrolateral sensorimotor cortex. Finally, we investigated Lumican expression in *Fezf2* null cortex, which completely and specifically lacks CSN and other SCPN (Chen et al., 2005; Molyneaux et al., 2005). Strikingly, Lumican expression is completely absent in the *Fezf2* null cortex (Fig. 1I), confirming by entirely independent approaches that Lumican is expressed essentially only by lateral SCPN.

We further and more broadly investigated Lumican expression throughout the central nervous system. In the postnatal forebrain, Lumican expression is highly restricted; in addition to SCPN in lateral cortex, Lumican is expressed by only a subset of caudoventral hippocampal cells (Fig. S1C). Lumican is also expressed by meningeal cells, encapsulating the central nervous system; this meningeal expression is seen at the level of the cortex (Fig. S1B–D), brainstem (Fig. S1E), and cervical spinal cord (Fig. S1F). Apart from this meningeal expression, Lumican is not expressed by any cells in the brainstem or cervical spinal cord. We also confirmed the specificity of the antibody used in these analyses by verifying that the expression observed in both cortex and meninges is abolished in *Lumican* null mice (Fig. S1B, D–F). Taken together, Lumican is highly specifically and transiently expressed by SCPN in lateral cortex during postnatal development, and is absent from adult SCPN.

The spatial restriction to only SCPN in lateral cortex strongly suggested that Lumican is specifically expressed by CSN_BC-lat_. We thus directly investigated whether Lumican is expressed by CSN_BC-lat_ by combining Lumican immunocytochemistry with retrograde analysis. We retrogradely labeled all CSN by injecting Cholera Toxin B subunit (CTB) into the cervical dorsal funiculus at P4, labeling all CSN in both medial and lateral cortex. Lumican is expressed by CSN in lateral cortex. Notably, there is striking and almost complete exclusion of Lumican from CSN in medial cortex (Fig. 1J). The fraction of Lumican-expressing, CTB-labeled CSN is significantly higher in lateral versus medial rostral sensorimotor cortex (40.0 ± 19.2% vs 7.7 ± 4.1%, respectively; mean ± SD; *p* < 0.05, unpaired two-tailed *t*-test; Fig. 1J). Together, these results identify that Lumican is expressed by CSN_BC-lat_ (Fig. 1K).

Proteoglycans, including Lumican and in particular including keratan-sulfate proteoglycans (KSPGs), are primarily ECM proteins, many of which have known functions in regulating axon pathfinding during development, and limiting plasticity following central nervous system injury (Caterson and Melrose, 2018; Ito et al., 2010; Jones and Tuszynski, 2002; Lowery and Van Vactor, 2009; Silver and Miller, 2004; Tessier-Lavigne and Goodman, 1996). Lumican core protein (predicted molecular weight; 37 kDa) is known to be post-translationally modified by N-linked glycans, which in certain instances are further modified by the addition of keratan sulfate glycosaminoglycan (GAG) chains to exist as a KSPG (Brézillon *et al*., 2013; Caterson and Melrose, 2018; Chen and Birk, 2013; Grover et al., 1995).

We therefore investigated whether Lumican expressed in the central nervous system undergoes either of these modifications. We dissected lateral sensorimotor cortex and cervical spinal cord from wild-type and *Lumican* null mice, followed by careful removal of meningeal membranes (Fig. S1G), then examined Lumican expression in these tissues by western blotting (Fig. S1H). In wild-type cortex and spinal cord, Lumican is detected as a single, narrow band at approximately 42 kDa, which likely represents the core protein. We also observed a broad band ranging from 55–75 kDa, presumably reflecting a mobility shift of modified Lumican protein likely corresponding to a glycoprotein/proteoglycan (Grover *et al*., 1995); both these bands are absent in tissue extracts from *Lumican* null mice (Fig. S1H). Since the molecular weight of Lumican in the central nervous system is lower than corneal Lumican (Fig. S1H), which contains highly sulfated GAG chains (Chakravarti et al., 1998), this result suggests that the GAG chains of Lumican in the central nervous system are presumably shorter or less sulfated compared to corneal Lumican. Importantly, we find that Lumican protein is efficiently secreted into the culture medium when overexpressed by cultured cortical primary neurons *in vitro* (Fig. S1I). These results indicate that CSN_BC-lat_-derived Lumican undergoes post-translational modifications, and that Lumican is efficiently secreted by neurons, consistent with previous findings identifying that Lumican is secreted by other cell types such as fibroblasts (Brézillon *et al*., 2013).

### Lumican suppresses CSN_medial_ axon collateralization in the cervical spinal cord, in a non-cell-autonomous manner

Lumican is expressed by CSN_BC-lat_ in the first two postnatal weeks, with peak expression occurring between P7 and P14 (Fig. 1 and S1). This occurs just after initial CSN axon segment-level targeting decisions have largely been made; CSN axons have already reached their target spinal segments in the CST (Bareyre et al., 2005; Kamiyama et al., 2015; Sahni *et al*., 2018), and are extending collaterals into the spinal gray matter (Kuang and Kalil, 1994). Thus, this timing of peak expression suggested that Lumican might likely control CSN axon collateralization rather than initial axon targeting.

Even more intriguingly, Lumican is expressed by only a highly selective CSN subpopulation—CSN_BC-lat_—that extend their axons only into bulbar and cervical spinal segments. The cervical spinal segments are also innervated by many CSN_medial_ (Sahni *et al*., 2018), which do not express Lumican (Fig. 1J). Like other known KSPGs, CSN_BC-lat_-derived Lumican is glycosylated (Fig. S1H). Further, cortical neurons secrete Lumican (Fig. S1I), suggesting that CSN_BC-lat_-derived Lumican might function in a non-cell-autonomous manner and regulate CSN_medial_ axon collateralization in the cervical spinal cord. Given the known inhibitory effects of KSPGs on axon growth and sprouting (Ito *et al*., 2010; Jones and Tuszynski, 2002), we hypothesized that Lumican might limit CSN_medial_ axon collateralization in the cervical spinal cord.

To test this hypothesis, we investigated CSN_medial_ axon collateralization in wild-type and *Lumican* null mice. To specifically analyze CSN_medial_ axon collateralization, we employed stereotactic injection of biotinylated dextran amine (BDA), an anterograde tracer, into caudomedial sensorimotor cortex of 4-week-old mice. One week after BDA injection, we analyzed BDA-labeled CSN_medial_ axons in the cervical cord (Fig. 2A). We first confirmed that BDA injection sites were in the appropriate medial location in sensorimotor cortex, and well-matched across animals, using three distinct criteria: 1) spread of BDA around the injection site in multiple series of coronal sections of the brain; 2) contralateral axon projections by BDA-labeled CPN; and 3) thalamic innervation by BDA-labeled CThPN neurons. We selected wild-type and *Lumican* null mice that had identical BDA injections (Fig. 2A–C and S2A) for further, unbiased and automated analysis of BDA labeled CSN_medial_ axon collateralization in the cervical cord, with blinded criteria established *a priori* (Fig. 2D–I and S2B, C).

**Figure 2:**
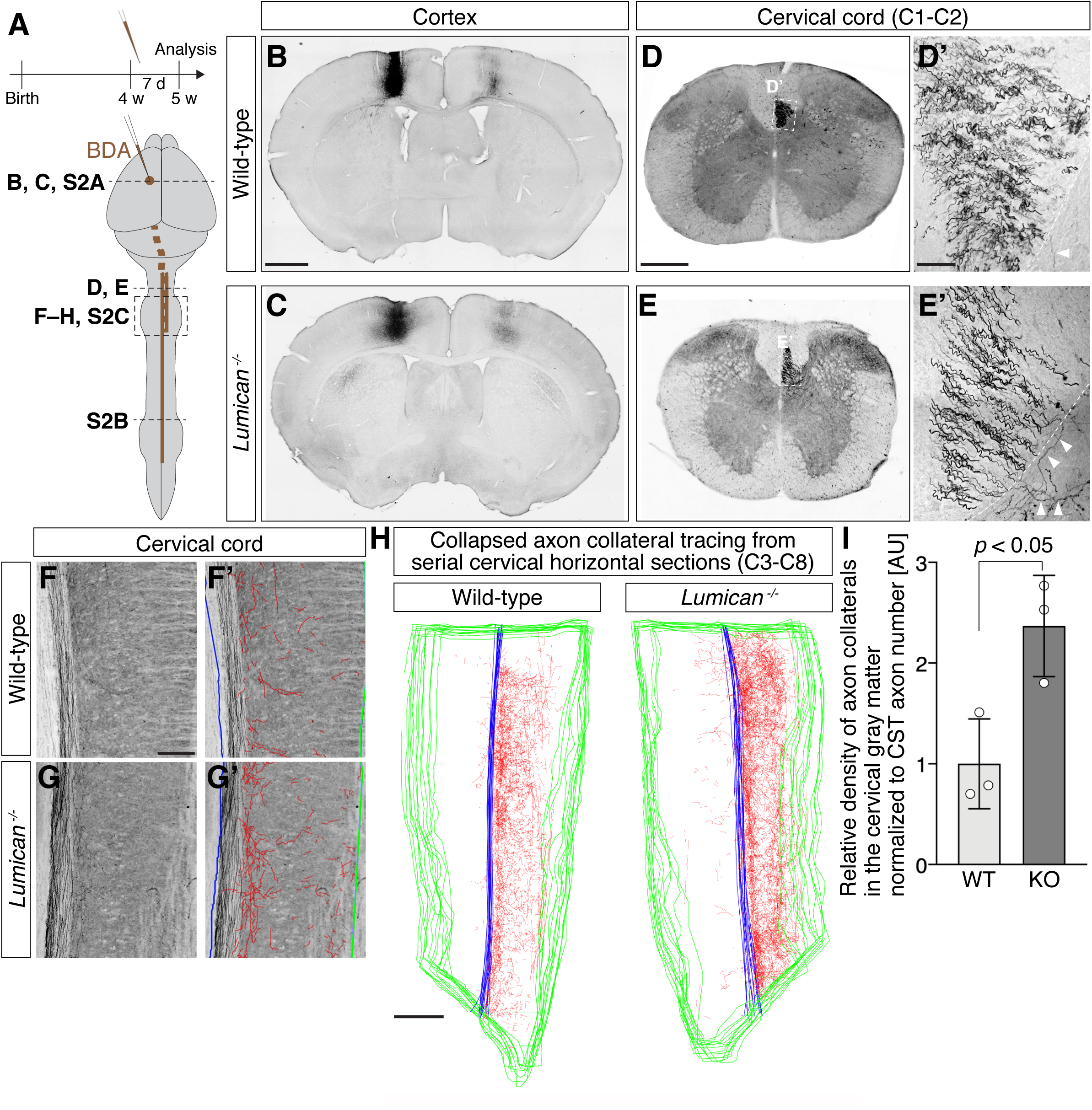
Lumican suppresses CSN_medial_ axon collateralization in the cervical spinal cord, in a non-cell-autonomous manner. (A) Schematic illustrating the experimental outline. (B, C) Representative coronal brain sections showing BDA injection site in M1. Successful BDA injection labels corticofugal projection through internal capsule, as well as callosal projection with homotopic contralateral specificity. BDA, black. Scale bar, 1 mm. (D, E) Axial sections of cervical cord (C1–C2 level) show corticospinal axons entering the spinal cord. CST axons in the dorsal funiculus are enlarged in magnified images (D’, E’). Note exuberant axon collateralization (white arrowheads) in *Lumican* null cervical gray matter, quite rare in wild-type. BDA, black. Scale bars = 500 µm (D, E); 50 µm (D’, E’). (F, G) Horizontal sections of cervical cord show collateral innervation into the gray matter. Manually traced axon collaterals are labeled in red. Midline and outer border of gray matter are labeled in blue and green, respectively (F’, G’). BDA, black. Scale bar = 200 µm. (H) Manually traced axon collaterals (red) on serial horizontal sections (C3–C8) are collapsed onto a single plane image, together with midline (blue) and gray matter border (green). Scale bar = 500 µm. (I) Quantification of relative corticospinal axon collateral density in the cervical gray matter normalized to CST axon number at C1–C2 shows increased CSN_medial_ axon collateralization in *Lumican* null mice compared to wild-type mice. Data are presented as mean ± SD, n = 3. WT, wild-type; KO, *Lumican* null.

Our experiments reveal that deletion of *Lumican* specifically de-represses CSN_medial_ collateralization in the cervical spinal cord. BDA labeled CSN_medial_ axons traversed the CST normally in the cervical dorsal funiculus in *Lumican* null mice, and a similar subset of CSN_medial_ axons reached the lumbar segment in both wild-type and *Lumican* null mice (Fig. 2D, E and S2B). In initial axial sections of the spinal cord at cervical C1–C2, there were visibly more exuberant axon collaterals in the cervical gray matter in *Lumican* null versus wild-type mice (Fig. 2D, E). Since CSN_medial_ axons extend widespread collaterals throughout the rostro-caudal extent of the cervical gray matter (Sahni *et al*., 2018), we next investigated CSN_medial_ axon collateralization across the entire cervical gray matter (from C3–C8). We used serial horizontal sections of the C3-C8 cervical spinal cord to visualize and quantify the total collateral area of BDA-labeled axons across the entire cervical spinal gray matter. Again, there is a dramatic increase in CSN_medial_ axon collateralization in *Lumican* null mice across the entire rostro-caudal extent of the cervical cord (Fig. 2F, G). We traced the total collateral area across the entire cervical gray matter (see Methods), and normalized this total area to the total number of labeled corticospinal axons in the dorsal funiculus at cervical C1–C2 to determine the approximate collateral target density in the cervical cord originating from each BDA labeled CSN_medial_ axon (Fig. 2D–I and S2C). This quantification shows increased CSN_medial_ axon collateral density in *Lumican* null mice compared to wild-type mice (Fig. 2I). Since CSN_medial_ do not express Lumican (Fig. 1), this result indicates that CSN_medial_ axon collateralization in the cervical spinal cord is suppressed by Lumican in a non-cell-autonomous manner.

We next investigated axon collateralization in the cervical gray matter by CSN_BC-lat_, which normally themselves express Lumican (Fig. S2D). Wild-type CSN_BC-lat_ axon collaterals are most abundant in the intermediate gray matter dorso-ventrally at cervical C1–C2 (Sahni *et al*., 2018) (Fig. S2D). In striking contrast to CSN_medial_, the density of cervical axon collaterals by CSN_BC-lat_ in *Lumican* null mice does not show a significant change when compared to wild-type; rather, though not significant, CSN_BC-lat_ axons even display a trend toward modest reduction in their cervical spinal collateral density (Fig. S2D; also see Discussion).

Taken together, these findings demonstrate that Lumican, produced by CSN_BC-lat_, non-cell-autonomously limits and relatively excludes CSN_medial_ axon collateralization in the cervical cord.

### CSN specification and main axon extension is unchanged in *Lumican* null mice

Given the highly specific Lumican expression by CSN_BC-lat_ in the postnatal central nervous system, we hypothesized that increased CSN_medial_ axon collateralization in *Lumican* null mice is specifically mediated by loss of Lumican expression by CSN_BC-lat_ rather than by some theoretically broader, non-specific defect in cortical organization. To rigorously test this hypothesis, we investigated earlier stages of cortical projection neuron specification in *Lumican* null mice.

We first examined expression of molecular controls over projection neuron subtype identity to investigate whether specification of broad cortical projection neuron subtypes might be altered in *Lumican* null cortex. Wild-type and *Lumican* null mice are indistinguishable in their overall brain size, or in their expression and laminar organization of CTIP2, SATB2, and Tbr1, a CThPN marker and control gene (Fig. 3A), confirming that Lumican does not function in early cortical development or cortical projection neuron specification.

**Figure 3:**
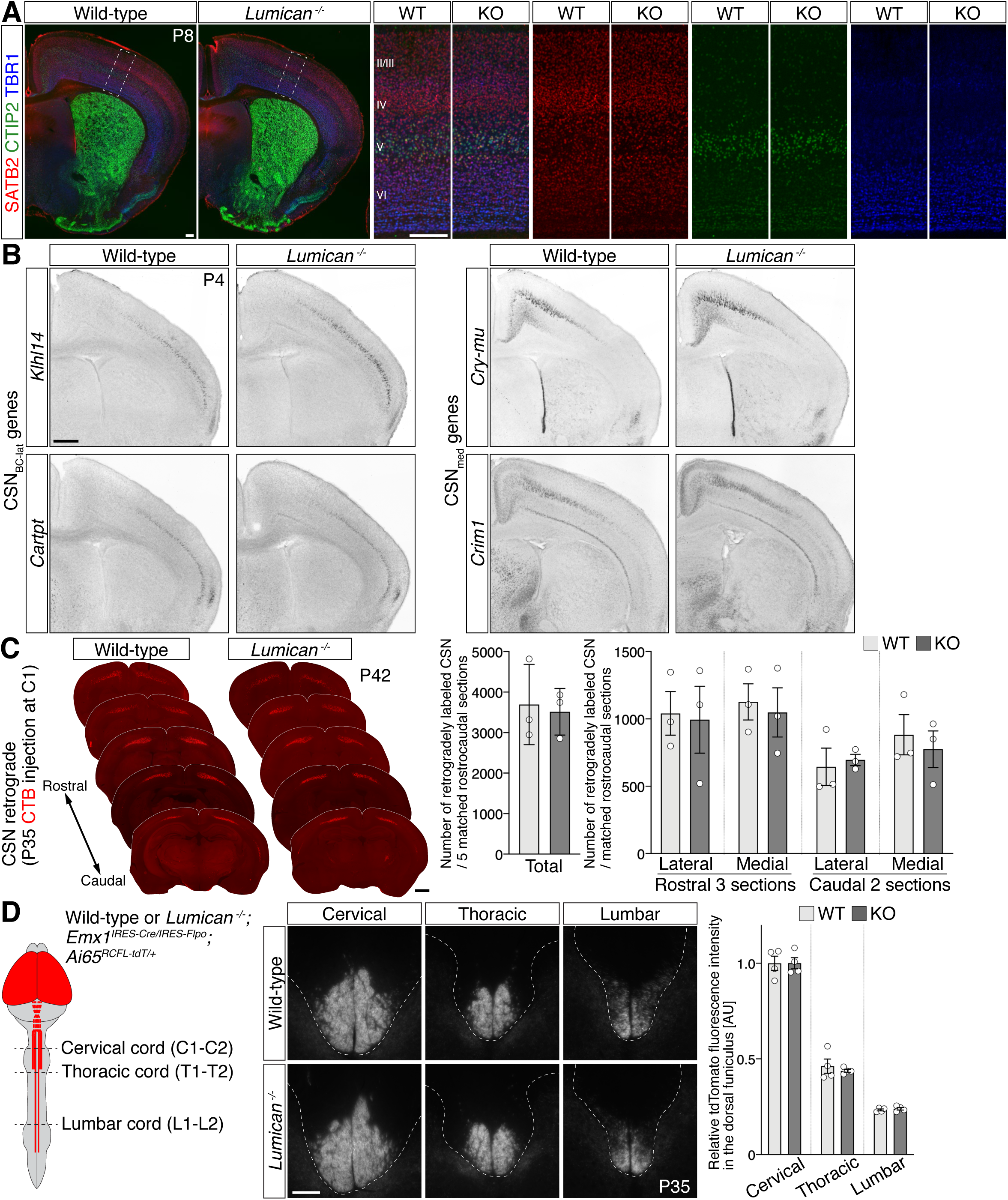
CSN specification and axon extension are unchanged in *Lumican* null mice. (A) In the absence of *Lumican*, overall brain structure remains unchanged. SCPN, CPN, and CThPN marker and control molecules CTIP2, SATB2, and TBR1, respectively, are expressed and positioned normally at P8. DAPI, blue. Scale bars = 200 µm. (B) Expression of CSN_BC-lat_-specific genes *Klhl14* and *Cartpt*, and CSN_medial_-specific genes *Cry-mu* and *Crim1* are essentially indistinguishable between wild-type and *Lumican* null cortex at P4. Scale bar = 200 µm. (C) CTB deposition at the top level of cervical segment (C1 level) at P35, followed by quantification of 5 matched rostro-caudal sections at P42 shows equivalent total number and distribution of retrogradely labeled CSN in wild-type and *Lumican* null cortex. Data are presented as mean ± SEM, n = 3. Scale bar, 1 mm. (D) CST axons in the dorsal funiculus are anterogradely labeled by *Emx1^IRES-Cre/IRES-Flpo^*;*Ai65^RCFL-tdT/+^.* Measurement of tdTomato fluorescence intensity in the dorsal funiculus white matter shows indistinguishable levels of signal at all spinal levels examined, and identical rostral-to-caudal reduction of tdTomato signal in wild-type and *Lumican* null mice, reflecting normal CST axon extension in *Lumican* null mice. Data are presented mean ± SEM, n = 4. Scale bar, 100 µm.

We next examined expression of genes normally expressed by distinct CSN subpopulations that we identified in recent work (Sahni *et al*., 2018) to investigate whether *Lumican* might control specification of these CSN subpopulations with segmentally distinct axonal projections in the spinal cord. CSN_BC-lat_ genes (*Klhl14* and *Cartpt*) and CSN_medial_ genes (*Crymu* and *Crim1*) are specifically expressed laterally and medially, respectively, in cortical layer V at P4 in wild-type mice, and their expression remains unchanged in *Lumican* null mice (Fig. 3B). Together, these data indicate that CSN are generated and specified normally in the absence of *Lumican*.

We next investigated the theoretical possibility that abnormal CSN_medial_ axon collateralization in *Lumican* null mice arises as a secondary consequence of abnormalities of CST axon segmental targeting. We directly investigated CSN axon extension in wild-type and *Lumican* null mice. Injection of CTB at the most rostral level of cervical segment (C1–C2 level) retrogradely labels all CSN. Both wild-type and *Lumican* null mice have efficient retrograde labeling, exhibiting indistinguishable total numbers and distribution of retrogradely labeled CSN (Fig. 3C), indicating that CSN axon extension into the spinal cord occurs normally in *Lumican* null mice. We also investigated CST axon extension using a genetic reporter; we crossed *Lumican* null mice with *Emx1^IRES-Cre/IRES-Flpo^*;*Ai65^RCFL-tdT/+^* mice, which express tdTomato in all cortical projection neurons (Sahni *et al*., 2018). CSN are the only cortical neurons that extend axons to the spinal cord, so this approach enables specific anterograde labeling of only corticospinal axons in the spinal cord. All wild-type CST axons extend to cervical C1–C2, then only a progressively diminishing subset of these axons extend to thoracic T1–T2 and lumbar L1–L2, as evidenced by progressive reduction of tdTomato intensity rostro-caudally, consistent with prior work (Bareyre *et al*., 2005; Sahni *et al*., 2018). This rostro-caudal distribution of CST main axon projection remains unaffected in *Lumican* null mice (Fig. 3D), indicating that Lumican does not control CST main axon extension. These data demonstrate that corticospinal axons reach and extend normally in the CST in *Lumican* null spinal cord.

Collectively, these results indicate that CSN specification and main axon extension occur normally in the absence of *Lumican*. Importantly, these results highlight that developmental processes taking place prior to corticospinal axon collateralization are essentially normal in *Lumican* null mice, indicating that the specificity of Lumican’s effect on CSN_medial_ axon collateralization does not arise as a secondary consequence of abnormalities in CSN specification or CST main axon segmental targeting.

### Lumican overexpression by CSN_BC-lat_ suppresses CSN_medial_ axon collateralization in the cervical cord

To investigate whether CSN_BC-lat_-derived Lumican directly suppresses CSN_medial_ axon collateralization, we next examined the effect of Lumican overexpression by CSN_BC-lat_ on CSN_medial_ axon collateralization. At P1 or P2, we injected AAV encoding either EGFP alone (AAV-EGFP) or EGFP and Lumican (AAV-EGFP-2A-Lumican) rostrolaterally to specifically target CSN_BC-lat_ in wild-type mice. We then used iontophoresis to stereotactically deliver BDA into caudomedial sensorimotor cortex in 4-week-old mice to analyze CSN_medial_ axon collateralization in the cervical cord (Fig. 4A). We applied criteria that were established blinded *a priori* to include only mice with precise and accurate AAV and BDA labeling for subsequent unbiased and automated analysis of axon collateralization. We rigorously ensured that there was no overlap between EGFP and BDA at all rostro-caudal levels of the cortex (Fig. S3A); that EGFP-labeled axons extended only to cervical and not to thoracic cord (Fig. 4B, C and S3A); and that BDA-labeled CSN_medial_ axons extended to thoracic cord and beyond (Fig. 4D, E and S3A). Together, these analyses confirmed that AAV and BDA labeling was specific to CSN_BC-lat_ and to CSN_medial_, respectively. We observed the expected level of previously described BDA-labeled CSN_medial_ axon collaterals across the rostro-caudal extent of the cervical spinal cord in control AAV-EGFP rostrolaterally injected mice (Fig 4D, F). In striking contrast, Lumican overexpression by CSN_BC-lat_ dramatically reduced collateralization in the cervical cord (Fig. 4E, G). Quantification of the relative collateral density of labeled CSN_medial_ axons shows substantial reduction in AAV-EGFP-2A-Lumican versus AAV-EGFP injected mice (Fig. 4H). This result strongly reinforces that CSN_BC-lat_-derived Lumican suppresses CSN_medial_ axon collateralization in a non-cell-autonomous manner.

**Figure 4:**
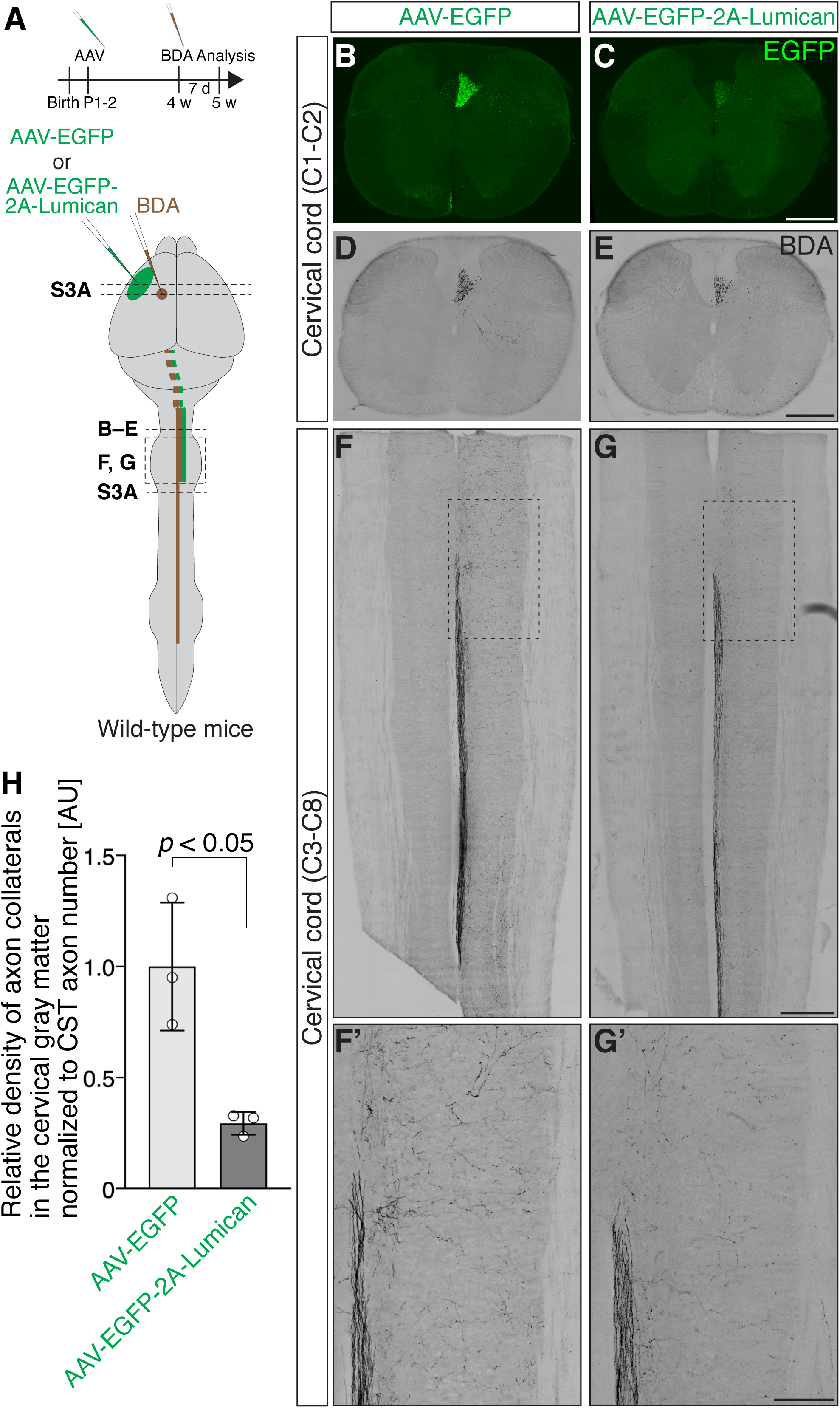
Lumican overexpression by CSN_BC-lat_ non-cell-autonomously suppresses CSN_medial_ axon collateralization in the cervical spinal cord. (A) Schematic illustrating the experimental outline. (B–E) Axial sections of C1–C2 cervical cord show corticospinal axons labeled by AAV-derived EGFP (green; B and C) or BDA (black; D and E). Scale bars = 500 µm. (F, G) Horizontal sections of cervical cord (C3–C8) display CSN_medial_ axon collateral innervation into the gray matter. BDA, black. Scale bars = 500 µm (F, G); 200 µm (F’, G’). (H) Quantification of relative corticospinal axon collateral density normalized to CST axon number at C1–C2 shows that Lumican overexpression by CSN_BC-lat_ substantially reduces CSN_medial_ axon collateralization. Data are presented as mean ± SD, n = 3.

Given the CSN axon collateralization phenotype in cervical cord in *Lumican* null mice, without any discernible effect on CSN main axon extension (Fig. 3C, D), we hypothesized that Lumican expressed by CSN_BC-lat_ suppresses CSN_medial_ axon collateralization locally in the spinal cord. We first investigated whether Lumican is trafficked into CSN axons. We overexpressed Lumican rostrolaterally in cortex in wild-type mice, and find that, consistent with this hypothesis, Lumican protein is present within CSN_BC-lat_ axons in the cervical cord (Fig. S3B). We also identified that endogenous Lumican protein is present in the cervical cord (Fig. S1H), and that Lumican is efficiently secreted by primary neurons (Fig. S1I), together suggesting that Lumican is anterogradely trafficked in CSN_BC-lat_ axons and likely secreted by these axons in the cervical cord.

We further investigated whether Lumican mis-expression locally in the cervical spinal cord suppresses CSN_medial_ axon collateralization (Fig. S3C). AAV-EGFP or AAV-EGFP-2A-Lumican was injected into the cervical gray matter at P1, when most CSN axons have not yet innervated the spinal gray matter, thereby limiting Lumican overexpression specifically to local spinal cells, and avoiding AAV infection of CSN. We indeed find very few (<10/mouse) EGFP-labeled CST axons, indicating that Lumican mis-expression is almost entirely limited to local spinal cells. We next analyzed CSN_medial_ axon collateralization labeled by AAV encoding tdTomato delivered into caudomedial sensorimotor cortex at P4. We find that Lumican mis-expression in cervical spinal cord substantially decreases CSN_medial_ axon collateralization at P25 (Fig S3C), indicating that Lumican can function locally in cervical spinal cord to suppress CSN_medial_ axon collateralization. Together, these results strongly support the interpretation that CSN_BC-lat_-derived Lumican suppresses CSN_medial_ axon collaterals locally in the cervical spinal cord.

### Lumican suppresses both bulbar-cervical and thoraco-lumbar CSN_medial_ subpopulations: intersectional AAV labeling identifies that CSN_BC-med_ axon collateralization is increased in *Lumican* null cervical cord

We have thus far used spatial locations of CSN within sensorimotor cortex (rostrolateral vs. caudomedial) to investigate CSN axon collateralization. However, CSN_medial_ are recently known to comprise at least two distinct subpopulations— the more commonly considered CSN_TL_ that extend thoraco-lumbar projections, and developmentally and molecularly distinct CSN_BC-med_ that extend axons exclusively to bulbar-cervical segments (Fig. 5A) (Sahni *et al*., 2018). BDA or AAV injections into caudomedial sensorimotor cortex, as described earlier, labels both CSN_BC-med_ and CSN_TL_. Importantly, axons of both subpopulations extend collaterals into the cervical spinal gray matter, though by CSN_TL_ at much lower density (Sahni *et al*., 2018). We therefore next investigated whether Lumican regulates axon collateralization by only one or by both CSN_medial_ subpopulations. Since CSN_BC-med_ and CSN_TL_ reside in a spatially interdigitated manner in caudomedial sensorimotor cortex (Sahni *et al*., 2018), this necessitated more advanced approaches than conventional anterograde tracing to investigate axon collateralization by these two distinct subpopulations. We applied two independent approaches—intersectional viral labeling (Fig. 5B–E and S4), which uses their segmentally distinct axon projection, and intersectional mouse genetics (Fig. 6 and S5), which takes advantage of the fact that CSN_BC-med_ and CSN_TL_ are molecularly distinct—to distinguish the two subpopulations.

**Figure 5:**
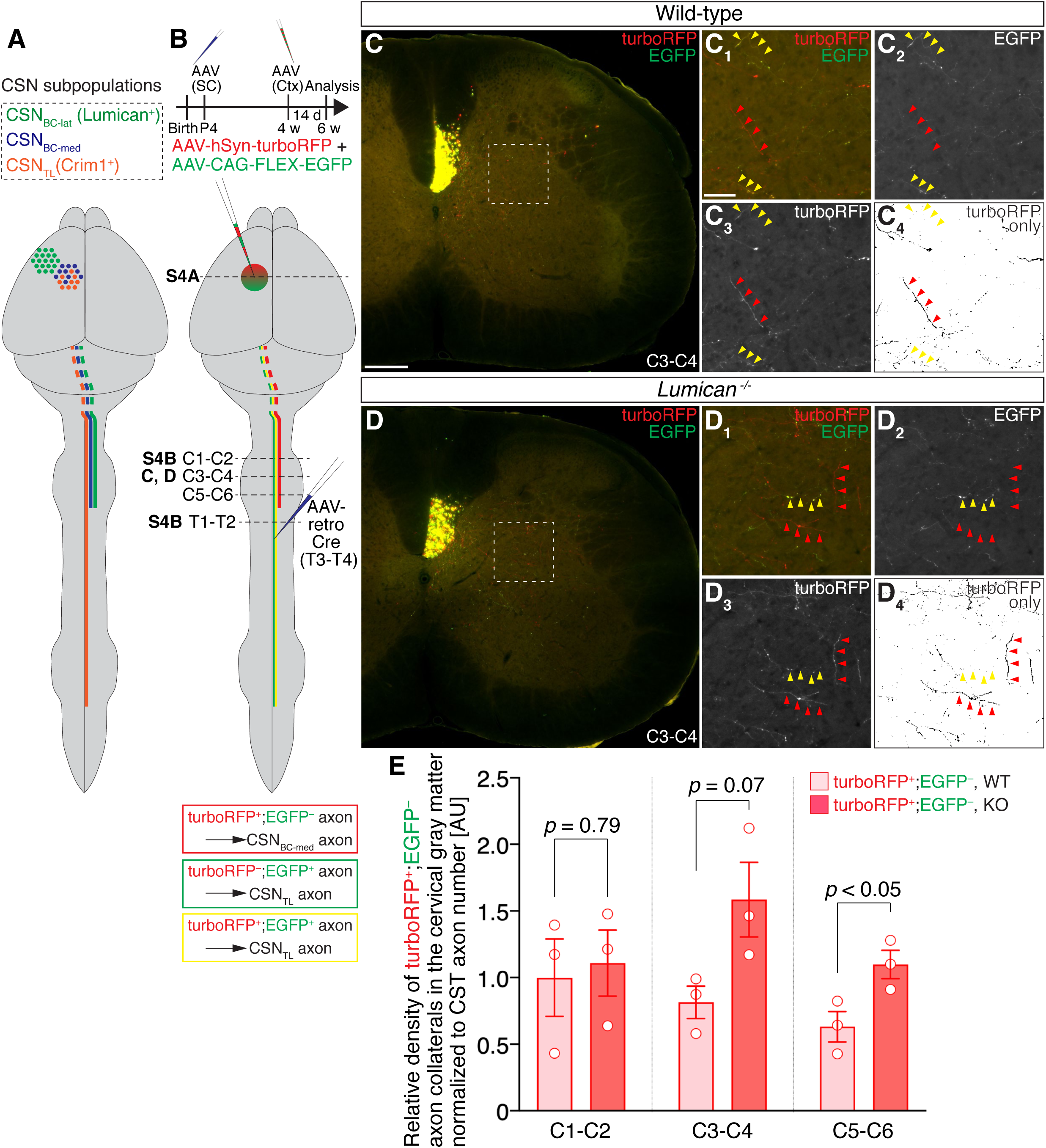
Exclusionary, subtractive viral labeling reveals increased CSN_BC-med_ axon collateralization in *Lumican* null cervical cord. (A) Schematized representations of developmentally distinct CSN subpopulations by the combination of location and projection specificity. Compared to the schematic in Fig. 1A, CSN in caudomedial sensorimotor cortex (CSN_medial_) are further subdivided into CSN_BC-med_ (blue) and CSN_TL_ (orange). While CSN_BC-med_ send projections only to brainstem and cervical cord, CSN_TL_ axons extend and collateralize throughout the spinal cord. For simplicity, axon collaterals are not illustrated, and CSN are illustrated in only one hemisphere. (B) Schematic illustrating the experimental outline. (C, D) Axial sections of C3–C4 cervical cord show turboRFP and EGFP fluorescence in the dorsal funiculus and gray matter. In higher-magnification images (C_1-4_, D_1-4_), red arrowheads indicate turboRFP^+^;EGFP^−^ axon collaterals, while yellow arrowheads indicate turboRFP^+^;EGFP^+^ axon collaterals. To identify turboRFP^+^;EGFP^−^ collateral signal, turboRFP^+^ and EGFP^+^ pixels above threshold are separately identified, followed by subtraction of EGFP^+^ pixels from turboRFP channel (C_4_, D_4_). Scale bars, 200 µm (C, D); 50 µm (C_1-4_, D_1-4_). (E) Quantification of relative density of turboRFP^+^;EGFP^−^ axon collateral density normalized to CST axon number at C1–C2. Data are presented as mean ± SEM, n = 3.

**Figure 6:**
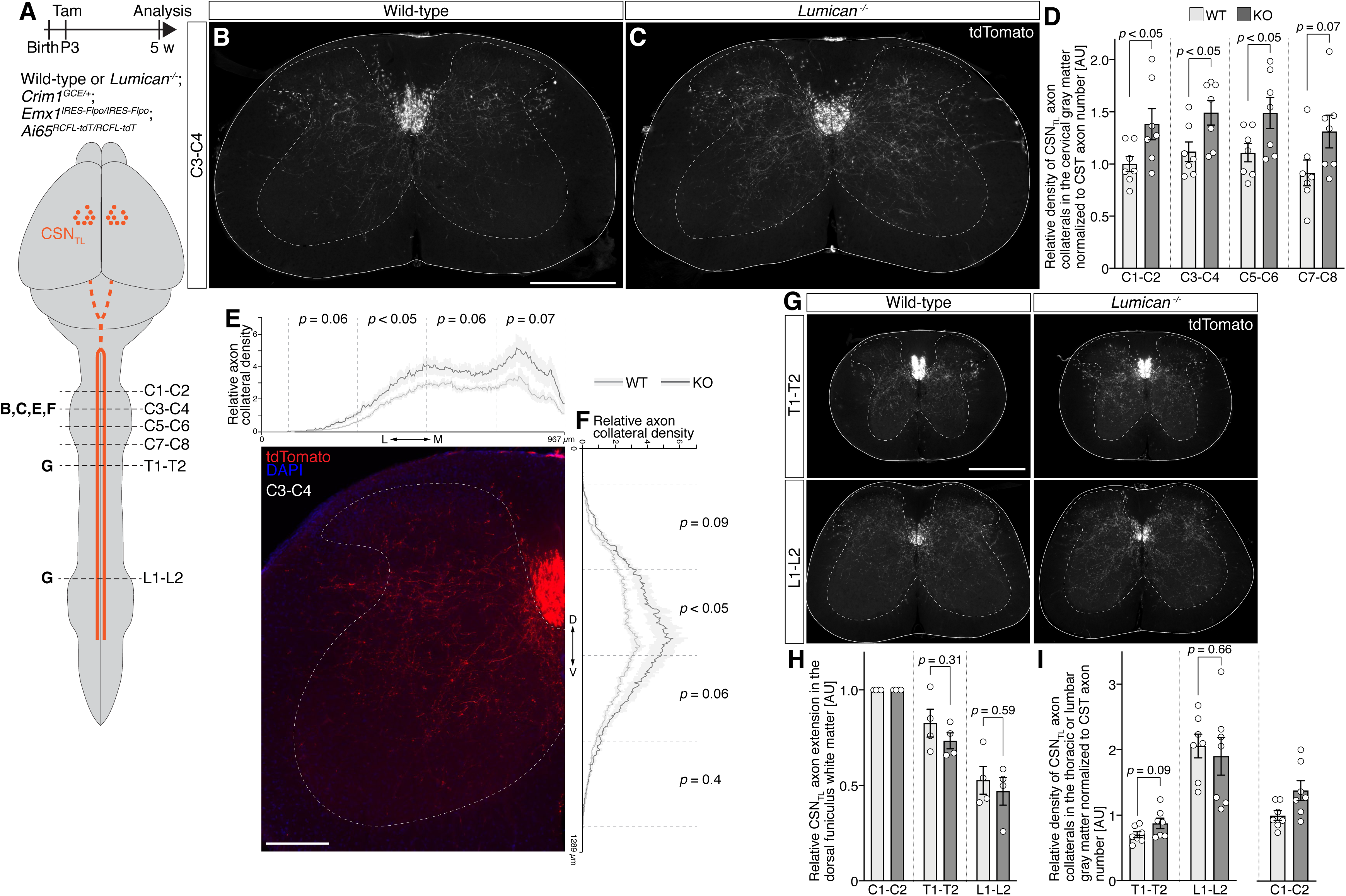
Intersectional genetic labeling reveals increased CSN_TL_ axon collateralization in *Lumican* null cervical cord. (A) Schematic illustrating the experimental outline. (B, C) Immunocytochemistry on axial sections of C3–C4 cervical cord at 5 weeks old display tdTomato^+^ corticospinal axons labeled by tamoxifen injection at P3 into *Lumican* wild-type or null *Crim1^GCE/+^;Emx1^IRES-Flpo/IRES-Flpo^;Ai65^RCFL-tdT/RCFL-tdT^* mice. Dashed lines demarcate gray matter. Scale bar, 500 µm. (D) Quantification of relative corticospinal axon collateral density normalized to CST axon number at C1–C2 shows increased CSN_TL_ axon collaterals in *Lumican* null mice compared to wild-type mice throughout the cervical cord. Data are presented as mean ± SEM, n = 7. (E, F) Distribution of CSN_TL_ axon collaterals was quantified along medio-lateral (E) and dorso-ventral (F) axes in wild-type and *Lumican* null cervical gray matter at C3–C4 level. A representative image is shown to delineate cervical gray matter (dashed line). Data are presented as mean ± SEM, n = 7. Statistical analysis was performed in each of four bins along each axis. Scale bar, 200 µm. (G) Immunocytochemistry on axial sections of T1–T2 thoracic cord and L1–L2 lumbar cord collected at 5 weeks old shows tdTomato^+^ corticospinal axons labeled by tamoxifen injection at P3. Dashed lines demarcate gray matter. Scale bar, 500 µm. (H) Quantification of CSN_TL_ axon numbers in the dorsal funiculus white matter shows indistinguishable rostral-to-caudal reduction in wild-type and *Lumican* null mice, reflecting normal CSN_TL_ main axon extension in *Lumican* null mice. Data are presented mean ± SEM, n = 4. (I) Quantification of relative CSN_TL_ axon collateral density at thoracic T1–T2 and lumbar L1–L2 normalized to CST axon number at each respective segment shows indistinguishable CSN_TL_ axon collateral density in *Lumican* null mice compared to wild-type mice. To enable direct comparison, the data for C1–C2 from (D) are reproduced here at this scale. Data are presented as mean ± SEM, n = 7.

We first investigated whether CSN_BC-med_ axon collateralization is regulated by Lumican. We utilized an intersectional viral labeling approach for exclusionary, subtractive labeling of the two CSN subpopulations within caudomedial sensorimotor cortex. This approach takes advantage of both the location of CSN_BC-med_ in caudomedial sensorimotor cortex, and the segmental projection of CSN_BC-med_ axons extending only to bulbar-cervical segments. When injected into caudomedial sensorimotor cortex, an AAV-expressed reporter under control of pan-neuronal promoter will label all neurons, including both CSN_TL_ and CSN_BC-med_, while a conditional, Cre-dependent AAV-expressed reporter combined with retrogradely transported AAV-Cre (AAV-retro-Cre(Tervo et al., 2016) injected into the thoracic cord will label only CSN_TL_, and not CSN_BC-med_. The combination of these two strategies enables delineation of CSN_BC-med_ as a specific subset within the spatially defined overall population of CSN_medial_. We first injected AAV-retro-Cre into thoracic T3–T4 in wild-type and *Lumican* null mice at P4. We then co-injected a Cre-dependent AAV-CAG-FLEX-EGFP along with AAV encoding turboRFP under the control of human *Synapsin 1* promoter (AAV-hSyn-turboRFP) into caudomedial sensorimotor cortex in 4-week-old mice. As schematized in Fig. 5B, CSN_TL_ are labeled by EGFP (tuboRFP^+^;EGFP^+^ or turboRFP^−^;EGFP^+^), while CSN_BC-med_ remain singly positive for turboRFP (turboRFP^+^;EGFP^−^).

Mice injected with all three AAVs showed turboRFP and EGFP expression around the injection site in medial cortex. As expected, turboRFP-positive neurons reside in both superficial and deep layers, whereas EGFP-positive neurons reside only in layer V in M1 (Fig. S4A). Within layer V in M1 around the injection site, this results in a highly interdigitated, “salt-and-pepper” positioning of the two intermingled populations—CSN_BC-med_ labeled only red, and CSN_TL_ labeled green or double labeled—as previously described (Sahni *et al*., 2018). As also expected, turboRFP-single-positive cells in layer V are more abundant in rostral sensorimotor cortex, while EGFP-positive cells are more abundant in caudal sensorimotor cortex (Fig. S4A), consistent with the gradual rostral-to-caudal shift from forelimb to hindlimb representations in motor cortex (Tennant *et al*., 2011). This topography is unchanged between wild-type and *Lumican* null mice (Fig. S4A).

We investigated axon extension by EGFP-positive and turboRFP-single-positive CST axons in the dorsal funiculus at cervical and thoracic spinal levels. At cervical C1–C2, both EGFP-positive and turboRFP-single-positive CSN axons appear evenly distributed and interdigitated, without any detectable somatotopic organization of CST axons between CSN_BC-med_ and CSN_TL_ (Fig. S4B). We next analyzed extension by these distinctly labeled CSN axons to thoracic spinal level in wild-type and *Lumican* null mice (Fig. S4C). On average, < 40% of turboRFP-single-positive CSN axons (CSN_BC-med_) at cervical C1–C2 extend even to thoracic T1–T2, indicating that the majority of these axons terminate within the cervical spinal cord. In contrast, ∼ 70% of EGFP-positive CSN axons at cervical C1–C2 extend to thoracic T1–T2. Together, this indicates that turboRFP-single-positive CSN axons are preferentially enriched for CSN_BC-med_, in contrast to EGFP-positive CSN axons, which are enriched for CSN_TL_. The percentage of turboRFP-single positive axons at cervical C1–C2 that extend to thoracic T1–T2 is indistinguishable between wild-type and *Lumican* null mice. This approach enabled investigation of CSN_BC-med_ axon collateralization as a distinctly labeled subset within the overall CSN_medial_ subpopulation.

We next quantitatively compared the target density of CSN_BC-med_ axon collaterals, labeled by only turboRFP, in cervical spinal gray matter between wild-type and *Lumican* null mice. Given the theoretical possibility that Lumican might control CSN_BC-med_ axon collateralization variably at distinct levels of the cervical spinal cord, we analyzed axon collateralization in axial sections of the cervical spinal cord at three distinct segments—C1–C2, C3–C4, and C5–C6. We find that CSN_BC-med_ axon collateral density is increased in the cervical spinal cord from C3 to C6 in *Lumican* null compared to wild-type mice (Fig. 5C–E). Interestingly, CSN_BC-med_ axon collateralization remains unchanged at cervical C1–C2 in *Lumican* null mice. These results indicate that CSN_BC-med_ axon collateralization in the cervical spinal cord is suppressed by Lumican.

### Lumican also suppresses the thoraco-lumbar CSN_medial_ subpopulation: intersectional genetic reporter labeling identifies that CSN_TL_ axon collateralization is increased in *Lumican* null cervical cord

Labeling a specific subset of neurons by mouse genetic tools provides a powerful, non-invasive approach for anatomical and functional investigation. The intersectional viral labeling approach described above enables enrichment of CSN_BC-med_, but CSN_TL_ axons labeled using this approach are axotomized by the AAV-retro-Cre injection in the thoracic dorsal funiculus at P4. This could potentially modify more rostral collateral formation, thus interfering with accurate investigation of CSN_TL_ axon collateralization in the cervical spinal cord. To circumvent this potential effect, we investigated CSN_TL_ axon collateralization using *Crim1^GCE^;Emx1^IRES-FlpO^;Ai65^RCFL-tdT^* intersectional genetic reporter mice (Sahni *et al*., 2018). *Crim1* expression during CSN development prospectively identifies CSN_TL_, and *Crim1^GCE^* knock-in mice express tamoxifen-inducible Cre recombinase (CreERT2) from the *Crim1* locus. Because there is some *Crim1* expression by neuronal populations in the spinal cord, including spinal motor neurons, these mice were bred with *Emx1^IRES-FlpO^* knock-in mice. In these mice, FlpO recombinase is expressed under the control of the *Emx1* locus, which is expressed by all neocortical projection neurons but not in the spinal cord. These mice were further bred with intersectional reporter *Ai65^RCFL-tdT^* mice, in which tdTomato reporter expression occurs only in cells that express both Cre and Flp recombinases (Madisen et al., 2015). In these triple transgenic mice, we previously validated that intersectional recombination occurs only in neocortex, and verified that tdTomato-labeled axons in the spinal cord are predominantly CSN_TL_ (Sahni *et al*., 2018).

To investigate CSN_TL_ axon collateralization in the absence or presence of Lumican function, we crossed these triple transgenic mice with *Lumican* null mice to generate *Lumican* _wild-type;*Crim1*_*_GCE/+_*_;*Emx1*_*_IRES-Flpo/IRES-Flpo_*_;*Ai65*_*_RCFL-tdT/RCFL-tdT_* _or *Lumican*_ null;*Crim1^GCE/+^*;*Emx1^IRES-Flpo/IRES-Flpo^*;*Ai65^RCFL-tdT/RCFL-tdT^* mice. Both sets of mice were injected with tamoxifen at P3 to label CSN_TL_ (Fig. 6A and S5). We analyzed axon collateralization in axial sections of the cervical spinal cord at distinct segments. As previously described (Sahni *et al*., 2018), CSN_TL_ axons in both wild-type and *Lumican* null mice extend collaterals in the cervical gray matter at all these spinal segments. We find that CSN_TL_ in *Lumican* null mice have more exuberant collaterals than CSN_TL_ in wild-type mice (Fig. 6B, C). Quantification of axon collateral density reveals increased axon collateralization by *Lumican* null CSN_TL_ throughout the cervical cord (Fig. 6D). This demonstrates that Lumican also suppresses CSN_TL_ axon collateralization throughout the cervical spinal cord.

We next investigated whether Lumican regulates the dorso-ventral and medio-lateral distribution of CSN_TL_ axon collateralization within the cervical cord. We used a binning analysis to quantify relative axon collateral density at cervical C3–C4 (Fig. 6E, F). In wild-type mice, CSN_TL_ axon collaterals are present predominantly in the cervical intermediate gray matter dorso-ventrally: ∼12% of CSN_TL_ axon collaterals are present in the dorsal-most quarter of the spinal gray matter; very few CSN_TL_ collaterals (∼7%) are present in the ventral-most quarter of the spinal gray matter; ∼81% of CSN_TL_ axon collaterals are present in the central half of the spinal gray matter (Fig. 6F). Further, ∼70% of CSN_TL_ axon collaterals are present in the medial half of the spinal gray matter, with very few (∼7%) collaterals present in the lateral most quarter of the spinal gray matter (Fig. 6 E). This overall topography of CSN_TL_ axon collateral distribution is unaltered in *Lumican* null mice (Fig. 6E, F). While proportionally increased throughout the medio-lateral extent of the cord (Fig. 6E), CSN_TL_ axon collateral density in *Lumican* null mice is relatively unchanged in either the dorsal or ventral gray matter (Fig. 6F). In contrast, CSN_TL_ collateral density is significantly increased throughout the intermediate gray matter in *Lumican* null mice (Fig. 6F), where axon collaterals of Lumican-expressing CSN_BC-lat_ are also located (Fig. S2D). These findings indicate that Lumican limits CSN_TL_ axon collateralization within its normal domain of the cervical gray matter.

Finally, given specific Lumican expression by CSN_BC-lat_ that extend axons exclusively to bulbar-cervical segments, we investigated whether Lumican suppresses CSN axon collateralization in a segment-specific manner, by analyzing CSN_TL_ axons at thoracic T1–T2 and lumbar L1–L2. We first investigated CSN_TL_ main axon extension in the dorsal funiculus white matter. Consistent with the results using *Emx1^IRES-Cre/IRES-Flpo^*;*Ai65^RCFL-tdT/+^* mice (Fig. 3D), there is no difference in CSN_TL_ main axon extension in the dorsal funiculus from C1–C2 to thoracic T1–T2, and to lumbar L1–L2 between *Lumican* wild-type and null mice (Fig. 6G, H). We next investigated CSN_TL_ axon collateralization. In wild-type mice, when compared to CSN_TL_ axon collateral target density at C1–C2, target density at T1–T2 is reduced by ∼30% (*p* < 0.01, unpaired two-tailed *t*-test), and is increased at L1–L2 by ∼2 fold (*p* < 0.001; Fig. 6D, I). Strikingly, neither CSN_TL_ axon collateral target density at T1–T2 nor at L1–L2 is distinguishable between *Lumican* wild-type and null (Fig. 6I). These findings indicate that Lumican suppresses CSN_TL_ axon collateralization specifically in cervical spinal cord, and does not regulate CSN_TL_ axon collateralization in the thoracic or lumbar cord.

Taken together, our results indicate by multiple independent approaches that Lumican is expressed selectively by CSN_BC-lat_, and suppresses both CSN_BC-med_ and CSN_TL_ axon collateralization in the cervical cord in a non-cell-autonomous manner.

## Discussion

Execution of skilled movement is enabled by precise connectivity between the cerebral cortex and its targets in the brainstem and spinal cord with segmental specificity, established during development via regulated CSN main axon extension, followed by collateralization. Cortical areal representation of forelimb/arm motor control has expanded greatly through mammalian evolution, highlighting cortical connectivity with the cervical spinal cord as particularly important and interesting. The cervical spinal cord is innervated in different ways, with varying levels of specificity and precision, by three anatomically and molecularly distinct CSN subpopulations— CSN_BC-lat_, CSN_BC-med_, and CSN_TL_ (Sahni *et al*., 2018).

In the work presented here, we first identified that the proteoglycan Lumican is expressed exclusively by CSN_BC-lat_. We further identified that Lumican non-cell-autonomously suppresses axon collateralization by both caudomedial CSN subpopulations— CSN_BC-med_ and CSN_TL_. It is noteworthy that CSN_BC-lat_, residing outside M1, control development of corticospinal axons arising from M1. Additionally, this suppression occurs independent of any effects on CSN main axon extension, indicating that molecular control over axon collateralization in the spinal gray matter is dissociable from control over CSN main axon extension in the spinal white matter (Kalil and Dent, 2014; Ozdinler and Macklis, 2006). These results identify an entirely novel, molecularly-mediated crosstalk between axons of distinct CSN subpopulations that refines their differential innervation of the cervical spinal cord during development.

We speculate that these findings might enable new insights and deeper understanding of how corticospinal circuitry for skilled motor control has evolved in mammals. We have recently characterized anatomical and molecular identity of CSN_BC-lat_ in mice and, through this recent and other work (Cederquist et al., 2013; Sahni *et al*., 2018), it seems likely that this CSN subpopulation is evolutionarily newer. We now identify a novel mechanism by which this (presumptively) evolutionarily newer subpopulation regulates axon collateralization by (presumptively) evolutionarily older CSN_medial_. It is intriguing to consider this as a form of “innervation competition”–that evolutionarily newer CSN collateralize and more effectively innervate targets for the finest motor control in the brainstem and cervical spinal cord at least in part by actively suppressing and thus rewiring connectivity of their evolutionarily older counterparts to less “critical” spinal cord targets. Notably, CSN_BC-lat_ in *Lumican* null mice exhibit a trend toward reduced axon collateralization (Fig. S2D), potentially reflecting a secondary effect of relatively increased axon collateralization by CSN_medial_ (Fig. 2). While it remains theoretically possible that Lumican also has cell-autonomous, autocrine function, which normally preserves CSN_BC-lat_ collateral space, the data from our experiments strongly suggest that CSN_medial_ axons have taken up CSN_BC-lat_ “collateral space” in *Lumican* null mice; CSN_medial_ appear to “compete” more effectively in the absence of Lumican.

Similar or related mechanisms might have likely emerged during mammalian evolution for refinement of corticospinal circuitry. A separate motor cortex emerged with placental mammals about 100 million years ago (Kaas, 2004). Non-primate early mammals then acquired additional motor areas, such as dorsal premotor and supplementary motor areas (Kaas, 2013). Interestingly, ventral premotor area is considered unique in primates (Kaas, 2013). Importantly, all these areas that appeared more recently through evolution also display corticospinal innervation of the cervical spinal cord in primates, suggesting the potential for inter-axonal crosstalk between these distinct corticospinal projections (Dum and Strick, 2002; Schieber, 2007).

Remarkably, expression analyses in the developing primate cortex reveal that SLRPs, including Lumican, are expressed in appropriate locations, and at the appropriate developmental times, potentially controlling corticospinal axon branching during development. Developing marmoset cortex at birth, whose neural development corresponds roughly to P10 in mice (Clancy et al., 2001), exhibits similar, but expanded, *Lumican* expression (Fig. S6A) (Bakola et al., 2015; Shimogori et al., 2018). Even more strikingly, another SLRP family member, *Decorin,* shows prominent layer V expression in ventrolateral cortex in marmosets (Fig. S6A) (Bakola *et al*., 2015; Shimogori *et al*., 2018), but not in mice (Fig. S6B, C) (Gong et al., 2003; Lein et al., 2007; Visel et al., 2004). Expanded use of SLRPs might underlie evolutionary rewiring of corticospinal circuitry, which could potentially underlie refinement of motor control.

Lumican regulates CSN axons at sites that are remote from their parental soma in a paracrine fashion, as is recently known to occur with a few other regulators, but Lumican’s mode of function is rather unique since Lumican suppresses axon collateralization by other populations of neurons without altering the trajectory of the main CST. Axon guidance is known to be mediated not only via attractive and repulsive cues provided by intermediate or final targets, but also via axon-axon interactions (Chédotal, 2019; Imai and Sakano, 2011; Kalil and Dent, 2014; Tessier-Lavigne and Goodman, 1996; Wang and Marquardt, 2013). Classically, axon-axon interactions are known to be mediated by direct contact of cell-surface molecules that induce adhesion or repulsion, e.g. leading to axon fasciculation or sorting. However, recent findings also illustrate the importance of secreted molecules in such axon-axon interactions (Imai and Sakano, 2011; Wang and Marquardt, 2013). Notable among such mechanisms are Semaphorin-3F (Sema3F) and Sonic Hedgehog (Shh), expressed respectively by a subset of olfactory sensory neurons and retinal ganglion cells to repel axon trajectory from other subsets of the same neuronal subtype (Peng et al., 2018; Takeuchi et al., 2010). Future studies can identify whether Lumican specifically controls axon branching, subsequent arborization, and/or later pruning during development. Lumican function identified in the work presented here will help elucidate how axons with similar identity establish and maintain distinct target innervation, and likely mediate distinct circuit function(s).

While the mode of action of Lumican is revealed here as “crosstalk” from the Lumican-expressing CSN_BC-lat_ to suppress axon collateralization of both CSN_TL_ and CSN_BC-med_ subpopulations, the exact molecular mechanism(s) by which Lumican regulates their axon collateralization remains unclear. SLRPs, including Lumican, regulate collagen fibril assembly in both redundant and non-redundant manners, and their dysfunction leads to disorganized collagen fibrils, and associated defects most evident in connective tissues (Brézillon *et al*., 2013; Chen and Birk, 2013; Kalamajski and Oldberg, 2010). It is possible that altered collagen organization by Lumican deficiency changes molecular and physical properties of the ECM scaffold, thereby affecting axon development. However, the expression of collagen type I, whose fibrilization is regulated by Lumican (Chen and Birk, 2013), appears to be limited to meninges and blood vessels during brain development (Lein *et al*., 2007), suggesting that this is unlikely to be the mechanism controlling CSN axon collateralization; it is likely that other types of collagens or other collagen-like domain–containing molecules might be involved (Ricard-Blum, 2011). Lumican also binds to various secreted or cell surface molecules, and modulates multiple signaling pathways (Brézillon *et al*., 2013). Further, the LRR domain is a versatile protein-interaction domain, and recent studies indicate that neuronal LRR transmembrane proteins engage in diverse intercellular interactions that lead to synapse formation (de Wit and Ghosh, 2016). It is interesting to speculate that Lumican might suppress axon/synapse development by binding to transmembrane proteins on the axonal membrane. It could thereby function as a ligand to modulate intracellular signaling, and/or function as a competitor to block intercellular transmembrane-protein interactions. We have used a HEK cell surface binding assay (Kim et al., 2006) to test potential Lumican binding to select candidate transmembrane proteins, as listed in Supplementary Figure S7A, based on their: 1) higher expression by CSN_medial_ in our earlier transcriptomic analysis (Sahni *et al*., 2018); 2) previously known interaction with Lumican (Brézillon *et al*., 2013); and/or 3) possession of potential LRR-binding motif (de Wit and Ghosh, 2016; Seiradake et al., 2014). None of these tested candidates reveal significant binding to Lumican (Fig. S7A–C). We also investigated whether exogenously added Lumican can bind to neural tissues using an on-section binding assay (Feiner et al., 1997). We do not detect Lumican binding, apart from efficient interaction with meninges (Fig. S7D, E). Future investigation could further elucidate more exactly how Lumican regulates axon development.

The relevance of initial development of connectivity and circuitry for later disease vulnerability is supported by increasingly emerging evidence. It appears increasingly likely that subtle perturbations of early, circuit-specific developmental controls can lead to formation of subtly altered circuitry that is more vulnerable to disease in later life, whether by primary dysfunction or by less effective maintenance. For example, resequencing of candidate CSN developmental control genes identified in our previous work (Arlotta *et al*., 2005; Lai et al., 2008) has revealed novel candidate disease genes, including *Lumican*, in ALS (Daoud *et al*., 2011). A unique nonsynonymous *Lumican* variant (Leu199Pro) was identified in both sporadic and familial ALS patients, and this variant was predicted to be deleterious for its function (Daoud *et al*., 2011). We have tested a mutant mouse Lumican protein which harbors the corresponding L199P substitution by overexpression in primary neurons *in vitro*. Interestingly, the amount of secreted mouse Lumican L199P mutant is reduced when compared to wild-type Lumican protein (Fig. S8). Analysis of larger ALS cohorts, as well as generation of L199P knock-in mice, would enable further investigation into potential involvement of altered Lumican activity in ALS pathogenesis.

CSN exhibit substantial targeting and circuit diversity, and likely have even much greater molecular and functional diversity than described here, considering the tremendous areal, segmental, functional, and evolutionary diversity in the brainstem and spinal cord. Recent advances in single-cell and subcellular sequencing and proteomic technologies will enable further identification of CSN heterogeneity (Economo et al., 2018; Hatch et al., Accepted; Poulopoulos et al., 2019; Tasic et al., 2018). Identifying specific somatic and subcellular gene expression, RNA and protein localization, and local translational regulation signatures of increasingly more specific CSN subpopulations will enable molecular manipulations and circuit analyses of increasingly more specific circuit- and functionally-defined subsets of CSN. Elucidating dynamic, subtype-specific RNA and protein molecular machinery in subtype-specific growth cones *in vivo* (Poulopoulos *et al*., 2019), as well as postsynaptic target cell identity (Ueno *et al*., 2018) of progressively more delineated CSN subpopulations will contribute substantially to parsing and understanding corticospinal circuitry development, function, and evolution of seemingly “layered”, increasingly precise and advanced function-specific neuron populations. It is also possible that Lumican might regulate other descending pathways either directly or indirectly, including rubrospinal and reticulospinal pathways, as well as local circuits in the spinal cord. Intriguingly, *Lumican* is reported to be upregulated upon pyramidotomy in adult rat cervical cord (Bareyre *et al*., 2002). It is interesting to consider that Lumican manipulation after CST injury might enable rewiring with greater precision for enhanced recovery (Bareyre *et al*., 2002; Ghosh *et al*., 2010; Oudega and Perez, 2012). Future investigation will progressively deepen understanding of corticospinal development, functional organization, evolution, and selective vulnerability to degeneration, which together might enlighten and enable novel approaches for circuit regeneration and repair.

## Materials and Methods

### Mice

All mouse studies were approved by the Harvard University IACUC, and were performed in accordance with institutional and federal guidelines. Wild-type mice on a C57BL/6J or CD1 background were obtained from Charles River Laboratories (Wilmington, MA). The day of birth was designated as P0. *Lumican^+/–^* mice were generously provided by Dr. Shukti Chakravarti (Chakravarti *et al*., 1998). All *Lumican^+/–^* mice used in this study were maintained on a CD1 or CD1/C57BL/6J mixed background. *Fezf2^−/−^* and *Emx1^IRES-Cre/IRES-Cre^* mice were generated previously (Gorski et al., 2002; Hirata et al., 2004), and have been previously described (Molyneaux *et al*., 2005). *Crim1^GCE/+^* mice were obtained from Jackson Laboratories (Sahni *et al*., 2018). CreERT2 activity was induced at P3 by injecting 50 µl of Tamoxifen (Sigma, T5648) solution dissolved in corn oil (Sigma, C8267) at 8 mg/ml. *Emx1^IRES-FlpO/IRES-FlpO^* mice were generated previously (Sahni *et al*., 2018). *Ai65^RCFL-tdT/RCFL-tdT^* mice were obtained from Jackson Laboratories, and genotyped using their recommended protocol.

### DNA constructs

To generate pAAV-EGFP-2A-Stop (in which a stop codon was placed in frame 3’ to the EGFP coding sequence), the *EGFP-2A-Stop* coding sequence was cloned into an AAV shuttle plasmid (obtained from the Massachusetts General Hospital Virus Core) that contains the following elements flanked by AAV2 ITRs: a CMV/β-actin promoter to drive the expression of the gene of interest, followed by the woodchuck hepatitis virus post-transcriptional regulatory element (WPRE), an SV40 polyadenylation signal, and a bovine growth hormone polyadenylation signal. Mouse *Lumican* cDNA (Open Biosystems, 3585672) was inserted 3’ to the *EGFP* coding sequence; the two ORFs were separated by a T2A linker sequence in-frame to create a bicistronic expression vector pAAV-EGFP-2A-Lumican (wild-type). pAAV-EGFP-2A-Lumican L199P was generated by site-directed mutagenesis to introduce a Leu 199 to Pro mutation. A vector expressing fusion protein of the FLRT3 ectodomain and Fc domain of human IGHG1 (FLRT3-Fc) was provided by D. Comoletti. Nucleotides encoding the FLRT3 ectodomain were excised to generate a vector expressing only the Fc domain, or were replaced by mouse *Lumican* cDNA to generate a Lumican-Fc–expressing plasmid. Expression plasmids or cDNAs for genes listed in Fig. S7A were provided by E. Kim, A. Ghosh, J. de Wit, G. Miyoshi, A. Poulopoulos, or Y. Mukouyama, purchased from Addgene or Sino Biological, or described previously (Sahni *et al*., 2018).

### Retrograde labeling by CTB

Developmental CSN at P4 were retrogradely labeled bilaterally from cervical or thoracic spinal cord by injecting 161 nl of the retrograde label cholera toxin B subunit (CTB) conjugated to Alexa 647 (CTB-647, 1 mg/ml in phosphate buffer saline (PBS); Thermo Scientific, C34778) into each side of the midline guided by ultrasound backscatter microscopy (VisualSonics, Vevo 3100) via a pulled glass micropipette (Drummond Scientific, 3-000-203-G/X) with a digitally-controlled, volume-displacement nanojector (Nanoject II, Drummond Scientific). For these neonatal injections, pups were anesthetized under ice for 4 minutes. After injections, the pups were placed on a heating pad at 37°C for recovery. Pups were perfused for Lumican immunocytochemical analysis at P5.

CSN at P35 were retrogradely labeled bilaterally from cervical C1 segment by CTB-555 (2 mg/ml in PBS; Thermo Scientific, C22843). For these adult injections, mice were anesthetized under isoflurane. After laminectomy at the C1 vertebral segment, 322 nl of CTB-555 solution was injected into each side of the midline via a pulled glass micropipette with a Nanoject II. The skin was then sutured, and mice were placed on a heating pad at 37°C for recovery. Mice were subsequently perfused at P42 for analysis of CTB-labeled CSN in cortex.

### Preparation of AAV particles

AAV2/1 particles for AAV-EGFP and AAV-EGFP-2A-Lumican expression were generated at the Massachusetts General Hospital Virus Core using established protocols (Maguire et al., 2013). AAV8 hSyn-EGFP-Cre (described as “AAV-hSyn-EGFP-Cre”) was obtained from the vector core at the University of North Carolina at Chapel Hill. AAV Retrograde pmSyn1-EBFP-Cre (described as “AAV-retro-Cre”) and AAV1 pCAG-Flex-tdTomato-WPRE (described as “AAV-CAG-FLEX-tdTomato”) were obtained from Addgene. AAV1.hSyn.TurboRFP.WPRE.rBG (described as “AAV-hSyn-turboRFP”) and AAV1.CAG.Flex.eGFP.WPRE.bGH (described as “AAV-CAG-FLEX-EGFP”) were obtained from the vector core at the University of Pennsylvania.

All virus work was approved by the Harvard Committee on Microbiological Safety, and conducted according to institutional guidelines.

### Anterograde and retrograde labeling with AAV in early postnatal pups

For anterograde labeling of cortical neurons via AAV-mediated gene delivery, P1, P2, or P4 pups were anesthetized using hypothermia, for which pups were placed under ice for 2–4 minutes. The cortex was visualized via ultrasound backscatter microscopy (VisualSonics, Vevo 770 and 3100), then injected using a pulled glass micropipette attached to Nanoject II digitally-controlled volume injection system. The shapes of the brain, lateral ventricle, and hippocampus, along with skull surface markers, served as landmarks for these intracranial injections. For retrograde CSN labeling by AAV injection into the spinal cord, pups were similarly anesthetized, and the spinal cord was visualized using ultrasound backscatter microscopy. For these intraspinal (dorsal funiculus) injections, the size and shape of spinal segments, along with the midline, served as reproducible landmarks. After injections using the Nanoject II, pups were placed on a heating pad for recovery. CTB-647 (0.1 mg/ml; Invitrogen, C34778) was mixed with AAV-retro-Cre solution to visualize the injection sites in the spinal cord.

AAV titers and volumes used: AAV-EGFP, 9.4 x 10^12^ GC/ml, 115 nl (unilateral, Fig. S3C) or 161 nl (Fig. 4 and Fig. S3A); AAV-EGFP-2A-Lumican, 3.6 x 10^12^ GC/ml, 115 nl (unilateral, Fig. S3C) or 230 nl (Fig. 4 and S3A,B); AAV-retro-Cre, 5.4 x 10^12^ GC/ml, 115 nl (bilateral, Fig. 5 and S4); AAV1-CAG-FLEX-tdTomato, 7.7 x 10^12^ GC/ml, 46 nl (Fig. S3C); AAV-hSyn-EGFP-Cre, 8.1 x 10^11^ GC/ml, 46 nl (Fig. S3C).

### Anterograde labeling by tracer and AAV injections into juvenile (4-week-old) mice

To anterogradely label CSN in caudomedial sensorimotor cortex, we used 10,000 Da lysine-fixable biotinylated dextran-amine (BDA; Invitrogen, D1956), iontophoretically delivered into the appropriate cortical location. A small craniotomy was made over the left hemisphere of anesthetized 4-week-old mice positioned in a stereotactic frame (Stoelting). A pulled glass micropipette (∼80-µm–inner diameter tip) loaded with a 10% solution of BDA in PBS was stereotactically positioned at the following coordinates; anterior-posterior (AP) ± 0 mm; medio-lateral (ML) +1.0 mm; dorso-ventral (DV) +0.8 mm from pia. Using an Isolated Pulse Stimulator (A-M Systems, Model 2100), intermittent pulses of 8 µA current were delivered for 7 seconds, with an inter-pulse interval of 7 seconds, for a total duration of 20 min. The skin was sutured, and mice were placed on a heat pad for recovery. Mice were perfused 7 days later for subsequent analysis.

To anterogradely label CSN via AAV-mediated gene delivery, we performed unilateral AAV microinjections into the appropriate location in sensorimotor cortex. A small craniotomy was performed over the left hemisphere of anesthetized 4-week-old mice positioned in a stereotactic frame, and a pulled glass micropipette loaded with AAV was stereotactically positioned at the following coordinates: For rostrolateral sensorimotor cortex injection, AP +1.5 mm; ML +3.0 mm; DV +1.0 mm from pia; for caudomedial sensorimotor cortex injection, see above. Following injection of AAV particles, the skin was sutured, and mice were placed on a heat pad to recover. Mice were perfused 14 days later for cytochemical analysis.

AAV titers and volumes used: AAV-hSyn-turboRFP, 9.6 x 10^12^ GC/ml (Fig. 5 and S4), 92 nl; AAV-CAG-FLEX-EGFP, 2.33 x 10^13^ GC/ml, 92 nl (Fig. 5 and S4); AAV-CAG-FLEX-tdTomato, 7.7 x 10^12^ GC/ml, 230 nl (Fig. S2D); AAV-hSyn-EGFP-Cre, 8.1 x 10^11^ GC/ml, 230 nl (Fig. S2D).

### Immunocytochemistry and *in situ* hybridization

Mice were transcardially perfused with cold PBS followed by 4% paraformaldehyde (PFA) in PBS, and brains and spinal cords were dissected and postfixed in 4% PFA/PBS at 4°C overnight. Spinal cords were sectioned on a cryostat (Leica, CM 3050S) at 50 µm (all figures except Fig. 6 or Fig. S3B) or 70 µm (Fig. 6), or on a vibrating microtome (Leica) at 50 µm (Fig. S3B). Brain sections were collected on a cryostat at 50 µm thickness. Non-specific binding was blocked by incubating tissue and antibodies in 2% donkey serum (Millipore, S30-100ML)/0.3% BSA (Sigma, A3059-100G) in PBS with 0.3% Triton X-100 or 0.3% BSA in PBS with 0.3% Triton X-100. In most instances, tissue sections were incubated with primary antibodies at 4°C overnight. Thicker sections (70 µm) of the spinal cord were incubated with primary antibodies at 4°C for 2 days. Secondary antibodies were chosen from the Alexa series (Invitrogen), and used at a dilution of 1:500 for 3–4 hours at room temperature (RT). For DAPI staining, tissue was mounted in DAPI-Fluoromount-G (SouthernBiotech, 0100-20).

Primary antibodies and dilutions used: goat anti-Lumican (R&D Systems, AF2745, 1:200); rabbit anti-Lumican (Abcam, ab168348, 1:100); rat anti-CTIP2 (Abcam, ab18465, 1:2,000); mouse anti-SATB2 (Abcam, ab51502, 1:500); rabbit anti-TBR1 (Abcam, ab31940, 1:500); rabbit anti-GFP (Invitrogen, A11122, 1:1,000); rabbit anti-RFP (Rockland, 600-401-379, 1:500). For Lumican staining in Fig. S3B, sections were incubated in 0.1 M citric acid, pH 6.0, for 5 min at 95–98°C for antigen retrieval prior to standard staining protocol.

BDA was visualized using an ABC-HRP kit (VECTOR laboratories, PK-4000) and 3,3’-Diaminobenzidine (Sigma, D4418). After mounting on gelatin-coated glass slides, sections were dehydrated in ethanol, cleared in xylene, and mounted in DPX mountant (Sigma, 06522).

*In situ* hybridization was performed as previously described (Sahni *et al*., 2018). The 3’ untranslated region of *Lumican* cDNA (nucleotides 1517 to 2004, NM_008524) was used as a probe. See (Sahni *et al*., 2018) for *Klhl14, Cartpt, Cry-mu, and Crim1* probes.

### Primary cortical neuron culture

Dissociated neurons from dissected wild-type P0 cortices were nucleofected with pAAV-EGFP-2A-Stop, pAAV-EGFP-2A-Lumican wild-type, or pAAV-EGFP-2A-Lumican L199P plasmid, using an Amaxa Mouse Neuron Nucleofector kit (Lonza, VPG-1001) as described previously (Wuttke et al., 2018), and were cultured for 7 hours in Neurobasal Medium (Gibco, 21103049) supplemented with 5% fetal bovine serum (VWR, 97068-085), 0.25% GlutaMAX (Gibco, 35050061), 2% B27 (Invitrogen, 17504044), and 0.6% glucose (Sigma, G6152), then for 7 days in Neurobasal Medium (Gibco, 21103049) supplemented with 1% GlutaMAX (Gibco, 35050061) and 2% B27 (Invitrogen, 17504044) on poly-D-lysine–coated 6 well plates.

### Immunoblot analysis

Tissues were microdissected, and the meningeal membrane was removed from both rostrolateral sensorimotor cortex and cervical spinal cord. Microdissected tissue was lysed in a buffer containing 50 mM Tris-HCl (pH 7.5), 150 mM NaCl, 1.0% Triton X-100, 1 mM dithiothreitol, 1 mM EDTA, and Halt inhibitor cocktail (Thermo Scientific, 78440). Lysates were centrifuged at 20,000*g* at 4°C for 15 min, and the resulting supernatants were analyzed.

Primary cortical neurons were washed with PBS, and lysed with a cell lysis buffer containing 20 mM Tris-HCl (pH 7.5), 150 mM NaCl, 10 mM β-glycerophosphate, 5 mM EGTA, 1 mM Na_4_P_2_O_7_, 5 mM NaF, 0.5% Triton X-100, 1 mM Na_3_VO_4_, 1 mM dithiothreitol, 1 μg/ml aprotinin, and 1 μg/ml leupeptin. The lysate was centrifuged at 20,000*g* at 4°C for 15 min to separate supernatant (soluble fraction) and pellet (insoluble fraction containing nuclei). The pellet was dissolved in RIPA buffer (50 mM Tris-HCl (pH 8.0), 150 mM NaCl, 1.0% Triton X-100, 0.5% sodium deoxycholate, 0.1% SDS, 1 mM dithiothreitol, and protease inhibitors (1 μg/ml aprotinin and 1 μg/ml leupeptin). Conditioned medium was centrifuged at 960g at 4°C for 10 min, and the resulting supernatant was further centrifuged at 20,000g at 4°C for 10 min to remove debris (Fig. S8).

Following standard Tris-glycine SDS–PAGE, resolved proteins were electroblotted onto PVDF membranes using semi-dry transfer. Ponceau S was used for staining the total protein transferred to the membrane. Membranes were incubated with primary antibodies diluted in 5% BSA in TBS with 0.1% Tween-20 or in Can Get Signal buffer (Toyobo, NKB-201). The following primary antibodies were used for immunoblotting: mouse anti-β-actin (Sigma, A5441, 1:5,000); rabbit anti-GFP (Invitrogen, A11122, 1:1,000); goat anti-Lumican (R&D Systems, AF2745, 1:200). HRP-conjugated secondary antibodies (Abcam, ab98693 and ab6721; Santa Cruz, sc-2020) were used for ECL imaging. Immunoreactive bands were detected by chemiluminescence using SuperSignal West Pico PLUS (Thermo Scientific, 34580), which was visualized using a CCD camera imager (FluoroChemM, Protein Simple). Fiji (Schindelin et al., 2012) was used to measure band intensities.

### HEK 293T cell surface binding assay

HEK 293T cells were transfected with constructs encoding a candidate transmembrane protein of interest, Lumican, or Fc-fusion protein as shown in Fig. S7A–C using FuGene 6 (Promega). Two days after transfection, cells expressing the transmembrane protein were incubated with conditioned medium obtained from cells expressing Lumican or the Fc-fusion protein for 1 hour at RT. Cells were then fixed and immunolabeled using goat anti-Lumican (R&D Systems, AF2745, 1:200) and donkey anti-goat IgG Alexa555 (Invitrogen, A-21432, 1:500) for Lumican detection, or using goat anti-human IgG Alexa555 (Invitrogen, A-21433, 1:500) for Fc detection. Mouse anti-myc 9E10 (Sigma, M4439, 1:1,000) was used for labeling Myc-Ntng1/2. Non-specific binding was blocked by incubating cells and antibodies in 2% donkey serum/0.3% BSA in PBS with 0.3% Triton X-100. For DAPI staining, cells were mounted in DAPI-Fluoromount-G.

### Section binding assay

HEK 293T cells were transfected with constructs encoding Fc, Lumican-Fc, or FLRT3-Fc using FuGene 6. A day later, cells were washed with PBS, and were further grown in Opti-MEM (Gibco, 51985034) for 3 days. After this 3-day period, conditioned medium was diluted with Opti-MEM (as indicated in Figure S7D) before application onto tissue sections. The presence of secreted Fc Fusion protein in the medium was confirmed by immunoblotting with goat anti-human IgG antibody (Invitrogen, A-21433, 1:500).

Following euthanasia, P10 *Lumican* null mice were immediately perfused with cold PBS, followed by brain dissection. The brain was fresh frozen in liquid nitrogen, then embedded in OCT compound (Sakura Finetek). Twelve µm thick coronal sections obtained on a cryostat were mounted on glass slides (VWR, 48311-703), and treated as described (Feiner *et al*., 1997). Briefly, sections were immediately postfixed in precooled methanol at –20°C for 7 min, washed twice with PBS, blocked with PBS containing 10% fetal bovine serum (VWR, 97068-085) for 15 min, and incubated with Opti-MEM conditioned medium described above for 1 hour at RT. Then, sections were fixed in 4% PFA/PBS for 5 min at RT, washed three times with PBS, and incubated with goat anti-human IgG Alexa555 (Invitrogen, A-21433, 1:500) in 0.3% BSA/PBS for 30 min at RT. Sections were then washed three times with PBS, and mounted in DAPI-Fluoromount-G.

### Imaging and quantification

For epifluorescence microscopy, tissue sections and cells were imaged using either a Nikon Eclipse 90i or NiE microscope (Nikon Instruments) with a mounted CCD or sCMOS camera (ANDOR Technology), respectively. Z stacks were collapsed using the “Extended Depth of Focus” function on the NIS-Elements acquisition software (Nikon Instruments). Images were processed using ImageJ software (NIH), Fiji, or Adobe Photoshop. For confocal imaging, samples were imaged on an LSM 880 (Zeiss).

To quantify retrogradely labeled CSN distributed in medial versus lateral locations in postnatal cortex, images of 2 coronal brain sections at specific rostral levels 300 µm apart were divided into 5 medio-lateral bins spanning the width of one cortical hemisphere, and medial versus lateral distinction was achieved by combining the 2 medial bins as medial, and the 3 lateral bins as lateral for CSN counts. The medial 2 bins approximately match M1 and M2 areas. One cortical hemisphere was analyzed for each mouse, and every labeled neuron was manually counted in each section using the Cell Counter function in Fiji. To quantify retrogradely labeled CSN distributed in medio-lateral and rostro-caudal locations in adult cortex, 5 matched rostro-caudal coronal brain sections of both cortical hemispheres were analyzed. Each cortical hemisphere was divided into 5 medio-lateral bins, and the 2 medial and 3 lateral bins were combined, respectively, as medial vs. lateral for CSN counts.

To quantify BDA-labeled CST axons at cervical C1–C2, the dorsal funiculus was imaged in axial sections of the cervical spinal cord using a 40X oil immersion objective on the 90i. These Z-stacks were imported into ImageJ, and each labeled axon was manually counted using the entire series of images within each Z-stack. To quantify BDA-labeled CSN axon collaterals from cervical C3–C8, horizontal sections of the cervical spinal cord were imaged using a 10X objective on the 90i. BDA tracing results shown in Fig. 2F–H and S2C were performed manually using Adobe Photoshop. Individual traced images of serial horizontal sections of spinal cord that contained any axon collaterals were then overlaid using the midline and tissue edges as landmarks. BDA collaterals in the cervical gray matter were also semi-automatedly traced and quantified using Image J, as previously reported (Grider et al., 2006) with modifications. Briefly, the largest Hessian with the smoothing scale 2 was applied to images before a threshold was applied. Then, pixels above the threshold were measured to quantify the total area of BDA-positive axon collaterals. After combining tracing results across all horizontal sections spanning the entire cervical gray matter, the resulting total area of BDA-positive axon collaterals was normalized to CST axon number at C1–C2 to calculate relative density of BDA-positive axon collaterals. Both manual and semi-automated tracing approaches gave similar quantification results, validating the semi-automated tracing approach.

To investigate CST axon extension in the spinal cord (Fig. 3D), tdTomato fluorescence intensity in the dorsal funiculus labeled by *Emx1^IRES-Cre/IRES-Flpo^*;*Ai65^RCFL-tdT/+^* was measured in axial sections at distinct spinal levels (cervical C1–C2, thoracic T1–T2, and lumbar L1–L2) using Fiji.

Density of axon collaterals labeled by tdTomato (in *Lumican* wild-_type;*Crim1*_*^GCE/+^*_;*Emx1*_*^IRES-Flpo/IRES-Flpo^*_;*Ai65*_*^RCFL-tdT/RCFL-tdT^* _or *Lumican*_ null;*Crim1^GCE/+^*;*Emx1^IRES-Flpo/IRES-Flpo^*;*Ai65^RCFL-tdT/RCFL-tdT^* mice, or mice injected with tdTomato-expressing AAV) was quantified by first measuring the number of CST axons in the dorsal funiculus at cervical C1–C2, T1–T2, and L1–L2 using axial sections. To quantify the number of CST axons consistently and reproducibly using criteria established *a priori*, a threshold was applied to confocal images (40X or 63X objective) obtained from C1–C2, T1–T2, and L1–L2 axial sections, and the number of axon cross-sections, seen as bright spots of appropriate size on the axial sections, was measured to obtain an estimate of the number of labeled tdTomato^+^ CST axons. Then, axial sections of the spinal cord were imaged using a 10X objective on the NiE, and the total area of axon collaterals in the gray matter was measured using the semi-automated tracing approach described above (using Fiji) (Grider *et al*., 2006). The smallest Hessian with the smoothing scale 0.55 was applied to images obtained from axial sections before a threshold was applied. Then, pixels above threshold were quantified as the total area of tdTomato-positive axon collaterals, then normalized to CST axon number at C1–C2, T1–T2, or L1–L2 to calculate relative density of tdTomato-positive axon collaterals in the gray matter. To quantify the distribution of axon collateralization, entire axial sections from each spinal cord at cervical C3–C4 were divided using Fiji into 600 bins medio-laterally (300 bins in each spinal hemicord) and 400 bins dorso-ventrally. Each bin corresponds to 3.2 µm in width medio-laterally and in height dorso-ventrally.

An intersectional AAV labeling approach was used to specifically label CSN_BC-med_ (Fig. 5 and S4), by labeling neurons and their axons with three patterns of fluorescent protein expression, as shown in Fig. 5B. In order to specifically analyze axons labeled only by turboRFP, Fiji was used to subtract EGFP^+^ pixels from the corresponding turboRFP image, thereby identifying axons with only turboRFP signal (see Fig. 5C_1-4_ and D_1-4_). To count CST axons in the dorsal funiculus, a similar subtractive approach was used, by which EGFP^+^ pixels were first subtracted from the corresponding turboRFP confocal image (63X objective) obtained from C1–C2 axial sections. Then, a threshold was applied to both EGFP and the resulting turboRFP images, followed by particle measurement of bright spots of appropriate size to obtain an estimate of the number of EGFP^+^ and turboRFP^+^:EGFP^−^ CST axons, respectively. To quantify the total area of axon collaterals, the smallest Hessian with the smoothing scale 0.55 was applied to both EGFP and turboRFP images obtained from axial sections using a 10X objective on the NiE. Then, a threshold was separately applied to EGFP and turboRFP images, followed by binarization. Subsequently, EGFP^+^ pixels were subtracted from the corresponding binarized turboRFP image, and turboRFP^+^ pixels above threshold were quantified as the total area of turboRFP^+^:EGFP^−^ axon collaterals. This area was then normalized to the turboRFP^+^:EGFP^−^ CST axon number at C1–C2 to calculate the relative density of turboRFP^+^:EGFP^−^ axon collaterals in the cervical gray matter.

### Statistical analyses

Data were analyzed in Microsoft Excel or GraphPad Prism with two-tailed unpaired Student’s *t*-tests. A *p*-value of <0.05 was considered statistically significant. No statistical methods were used to pre-determine sample sizes.

## Acknowledgements

We thank T. Addison, K. Wang, M. Wettstein, and K. Yee for superb technical assistance; members of the Macklis laboratory for scientific discussions and helpful suggestions; the Harvard Center for Biological Imaging for infrastructure and support; A. Poulopoulos for helpful discussions and reagents; M. Hibi, S. Chakravarti, D. Comoletti, E. Kim, A. Ghosh, J. de Wit, G. Miyoshi, and Y. Mukouyama for generous sharing of mice and reagents. This work was supported by grants from the National Institutes of Health (R01s NS045523 and NS075672, with additional infrastructure supported by NS049553, NS104055, and DP1 NS106665), the ALS Association, the Travis Roy Foundation, and Massachusetts Department of Public Health Spinal Cord Injury Fund to J.D.M.. Y.I. was partially supported by the Uehara Memorial Foundation, the Kanae Foundation for the Promotion of Medical Science, the Murata Overseas Scholarship Foundation, and the DEARS Foundation. V.S. was partially supported by National Institutes of Health K12 fellowship (NTRAIN/NICHD K12HD093427) and the DEARS Foundation. S.J.S. was partially supported by National Institutes of Health predoctoral NRSA F31 NS063516. J.D.M. is an Allen Distinguished Investigator of the Paul G. Allen Frontiers Group.

## Author Contributions

Y.I., V.S., S.J.S., and J.D.M. designed research; Y.I., V.S., and S.J.S. performed research; Y.I., V.S., S.J.S., and J.D.M. analyzed data; and Y.I., V.S., and J.D.M. wrote and edited the manuscript.

---

**Supplementary Figure S1:**
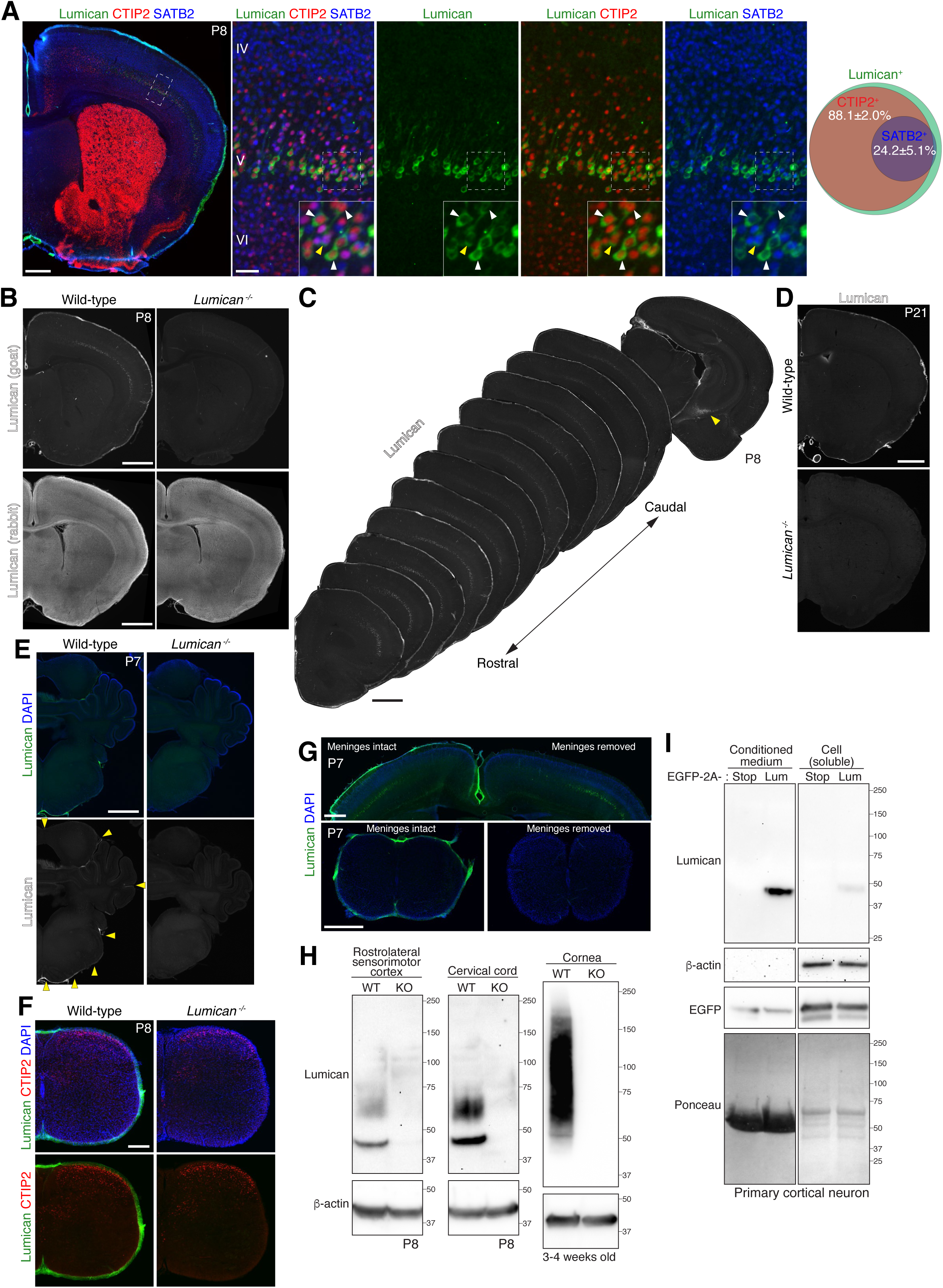
Lumican is expressed exclusively in the cortex within the central nervous system, and is secreted by primary cultured cortical neurons. (A) Immunocytochemistry on coronal section of P8 wild-type mouse brain showing Lumican (green), CTIP2 (red), and SATB2 (blue) expression. The overwhelming majority of Lumican-expressing cells are CTIP2^+^ (white arrowheads). A subset of Lumican^+^;CTIP2^+^ cells also express SATB2 (yellow arrowhead). Quantification is represented as Venn diagram (n = 3, P7 or P8, mean ± SEM). Note that almost all (95%, n = 2) SATB2^+^Lumican^+^ double-positive neurons also express CTIP2. DAPI, blue. Scale bars, 500 µm (low magnification); 50 µm (high magnification). (B) Lumican immunocytochemistry on coronal sections of P8 wild-type or *Lumican* null mouse brains. Lumican signal disappears in *Lumican* null brain, when detected with a highly specific goat anti-Lumican antibody. Although a single study reports Lumican expression in the cortex (Long *et al*., 2018), this work used a different antibody (rabbit anti-Lumican), which labels quite promiscuously and remains unchanged in *Lumican* null brain, suggesting lack of specificity of the rabbit antibody used in that study. Scale bars, 1 mm. (C) At P8, Lumican expression is strikingly specific to layer V across cortex, except caudomedial hippocampal cells (yellow arrowhead), and is prominently high rostrolaterally in the cortex. Scale bar, 1 mm. (D) Lumican is no longer expressed in cortex at P21. Scale bar, 1 mm. (E) Lumican is not expressed in the brainstem or cerebellum at P7. Yellow arrowheads indicate Lumican expression by meningeal cells. DAPI, blue. Scale bar, 1 mm. (F) In P8 cervical spinal cord, Lumican (green) is expressed by meningeal, but not by spinal cells. CTIP2, red. DAPI, blue. Scale bar, 200 µm. (G) Meningeal removal efficiently removes Lumican staining at the surface of mouse brain and spinal cord. In the upper panel, cortical meninges were removed from one hemisphere of P7 wild-type brain, and both hemispheres were processed for Lumican immunostaining (green). In the lower panels, P7 cervical cords with or without meningeal removal were immunolabeled for Lumican (green). Note that Lumican western blotting in Fig. S1H was conducted following meningeal removal. DAPI, blue. Scale bars, 500 µm. (H) Immunoblot analysis of P8 rostrolateral sensorimotor cortex, P8 cervical cord, and 3–4-week-old cornea in wild-type and *Lumican* null mice. In wild-type cortex and spinal cord, Lumican is detected as a single, narrow band at approximately 42 kDa, which likely represents the core protein, and as a broad band ranging from 55–75 kDa, presumably reflecting a mobility shift of modified Lumican protein likely corresponding to a glycoprotein/proteoglycan (Grover *et al*., 1995); both these bands are absent in tissue extracts from *Lumican* null mice. Tissue extracts from the cornea, where Lumican undergoes extensive KSPG modification (Chakravarti *et al*., 1998), exhibits a significant mobility shift of Lumican up to ∼150 kDa, as expected. (I) Immunoblot analysis of Lumican secretion in primary cortical neuron culture. Primary neurons derived from P0 mouse cortex were nucleofected with a plasmid encoding EGFP-2A-Stop or EGFP-2A-Lumican, and cultured for 7 days *in vitro*, followed by collection of conditioned medium and soluble cellular fraction. 1.9% and 24% of total conditioned medium and soluble cellular fraction was loaded, respectively. Note that most of the Lumican protein is secreted into the medium. Primary cortical neurons express Lumican core protein, but not the KSPG form. Conditioned medium and soluble cellular fraction samples were applied to the same gel, then the corresponding lanes were juxtaposed. Original gel images are shown in Supplementary Figure S8.

**Supplementary Figure S2:**
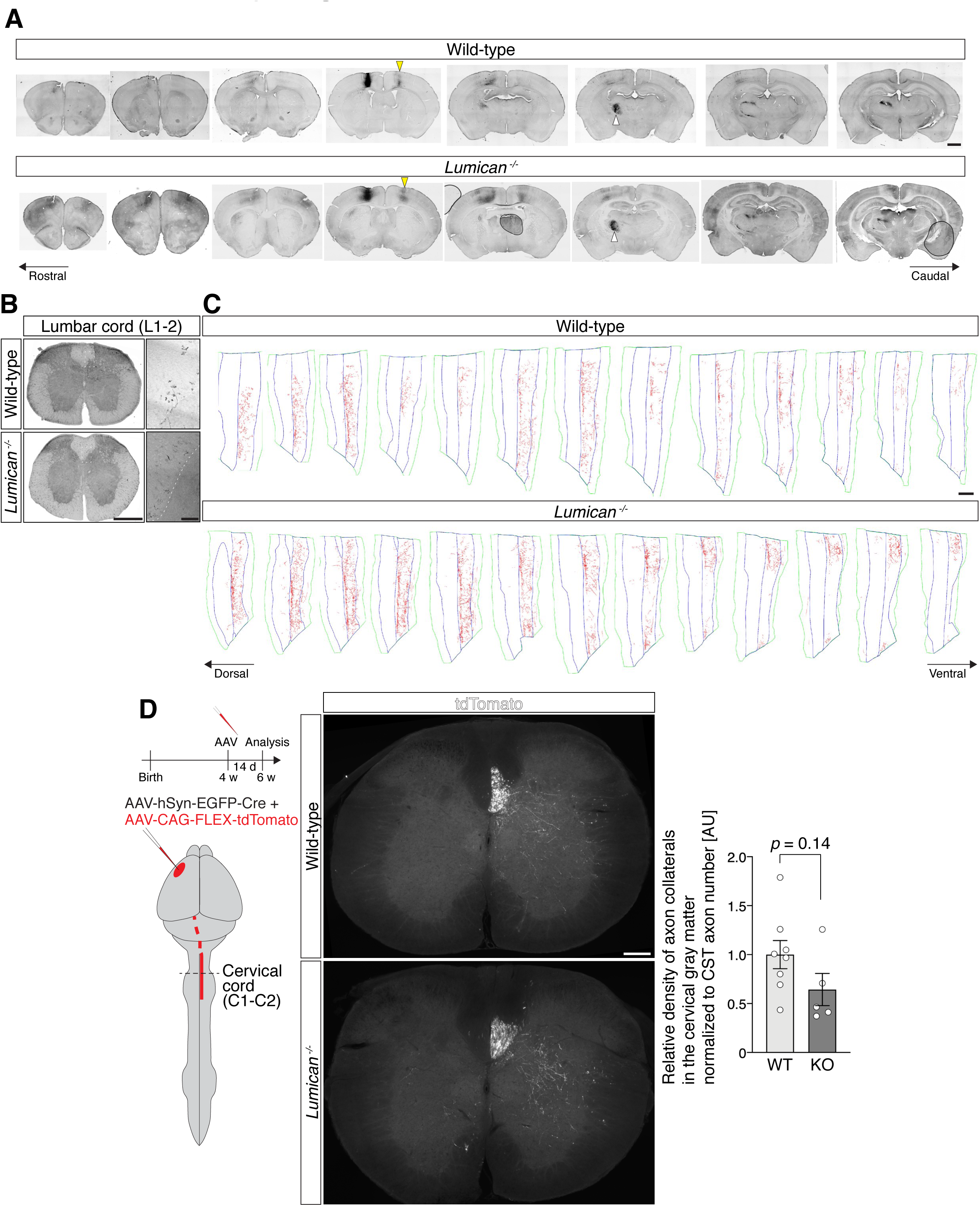
CSN_medial_ axon collaterals are increased, but CSN_BC-lat_ axon collaterals are not significantly changed, in *Lumican* null cervical cord. (A) Serial coronal brain sections showing BDA injection site in M1 and axon projections of BDA-labeled neurons. Matched BDA injections in wild-type and *Lumican* null brains exhibit comparable contralateral (yellow arrowhead) and thalamic (white arrowhead) innervation. Sections at the injection site are also shown in Fig. 2B, C. BDA, black. Scale bar, 1 mm. (B) Axial sections of lumbar cord (L1–L2 level) show corticospinal axons entering the lumbar cord. CST axons in the dorsal funiculus are enlarged in magnified images. BDA, black. Scale bars, 500 µm; 50 µm (inset). (C) Manually traced axon collaterals (red) on serial, individual horizontal sections (C3–C8) are shown. Midline and gray matter border, blue. Tissue border, green. All the sections spanning cervical gray matter (most are shown here) are projected onto one single plane as shown in Fig. 2H. Scale bar, 500 µm. (D) Schematic illustrates the experimental outline. AAVs encoding Cre recombinase (at a lower titer) and Cre-dependent tdTomato (at a higher titer) are stereotactically injected into rostrolateral sensorimotor cortex. Axial sections of C1–C2 cervical cord display tdTomato fluorescence in the dorsal funiculus and gray matter. Quantification of relative density of CSN_BC-lat_ axon collaterals normalized to CST axon number at C1–C2 does not show a significant change in *Lumican* null compared to wild-type mice, although there appears to be a trend toward a modest reduction in *Lumican* null mice (see Discussion). Data are presented as mean ± SEM, n = 8 (wild-type) or 5 (*Lumican* null). Scale bar, 200 µm.

**Supplementary Figure S3:**
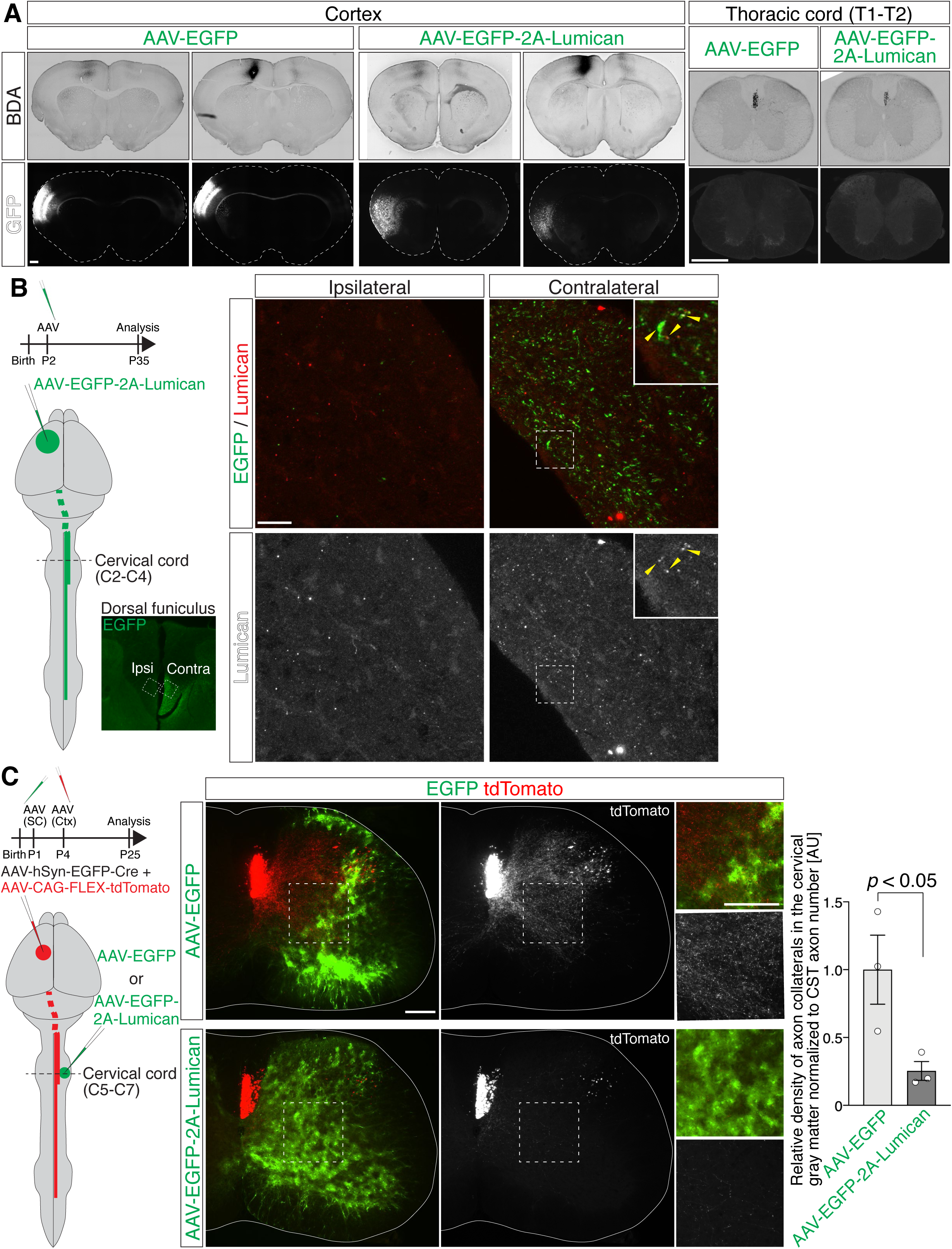
Lumican suppresses CSN_medial_ axon collateralization locally in the cervical spinal cord. (A) Coronal sections of 5-week-old wild-type brains injected with AAV encoding only EGFP or EGFP-2A-Lumican, in combination with BDA stereotactic injection (see Fig. 4A for experimental outline) demonstrate no overlap between EGFP and BDA staining, confirming entirely distinct and separated injections of AAV and BDA in cortex. Coronal sections around AAV and BDA injection sites are shown. Axial sections of thoracic cord at T1–T2 level exhibit abundant BDA, but very little EGFP signal in the dorsal funiculus and gray matter, while cervical cord exhibits both BDA and EGFP signals (Fig. 4B–E), confirming precise labeling of CSN_BC-lat_ and CSN_medial_ by AAV and BDA injections, respectively. Scale bars, 500 µm. (B) Lumican is trafficked to the spinal cord from the cortex. AAV virus encoding EGFP-2A-Lumican was injected into sensorimotor cortex of wild-type mouse brain at P2. Mice were fixed at P35, and axial sections of cervical spinal cord (C2–C4) were immunolabeled with antibodies against EGFP (green) and Lumican (red). Yellow arrowheads indicate colocalization of EGFP and Lumican in the contralateral dorsal funiculus. Scale bar, 10 µm. (C) Lumican mis-expression in the cervical cord suppresses CSN_medial_ axon collateralization. At P1, AAV encoding only EGFP or EGFP-2A-Lumican was injected into cervical gray matter. At P4, AAVs encoding Cre recombinase (at a lower titer) and Cre-dependent tdTomato (at a higher titer) were injected into caudomedial sensorimotor cortex to label CSN_medial_. Axial sections of C5–C7 cervical cord show EGFP (green) and tdTomato (red) fluorescence in the gray matter. Note that there are very few EGFP^+^ axons in the dorsal funiculus. Mis-expression of Lumican substantially reduces the amount of tdTomato^+^ CSN_medial_ axon collateralization. Quantification of relative density of axon collaterals normalized to CST axon number at C1–C2 shows substantial suppression of CSN_medial_ axon collateralization by Lumican mis-expression in the cervical cord. Data are presented as mean ± SEM, n = 3. Scale bars, 200 µm.

**Supplementary Figure S4:**
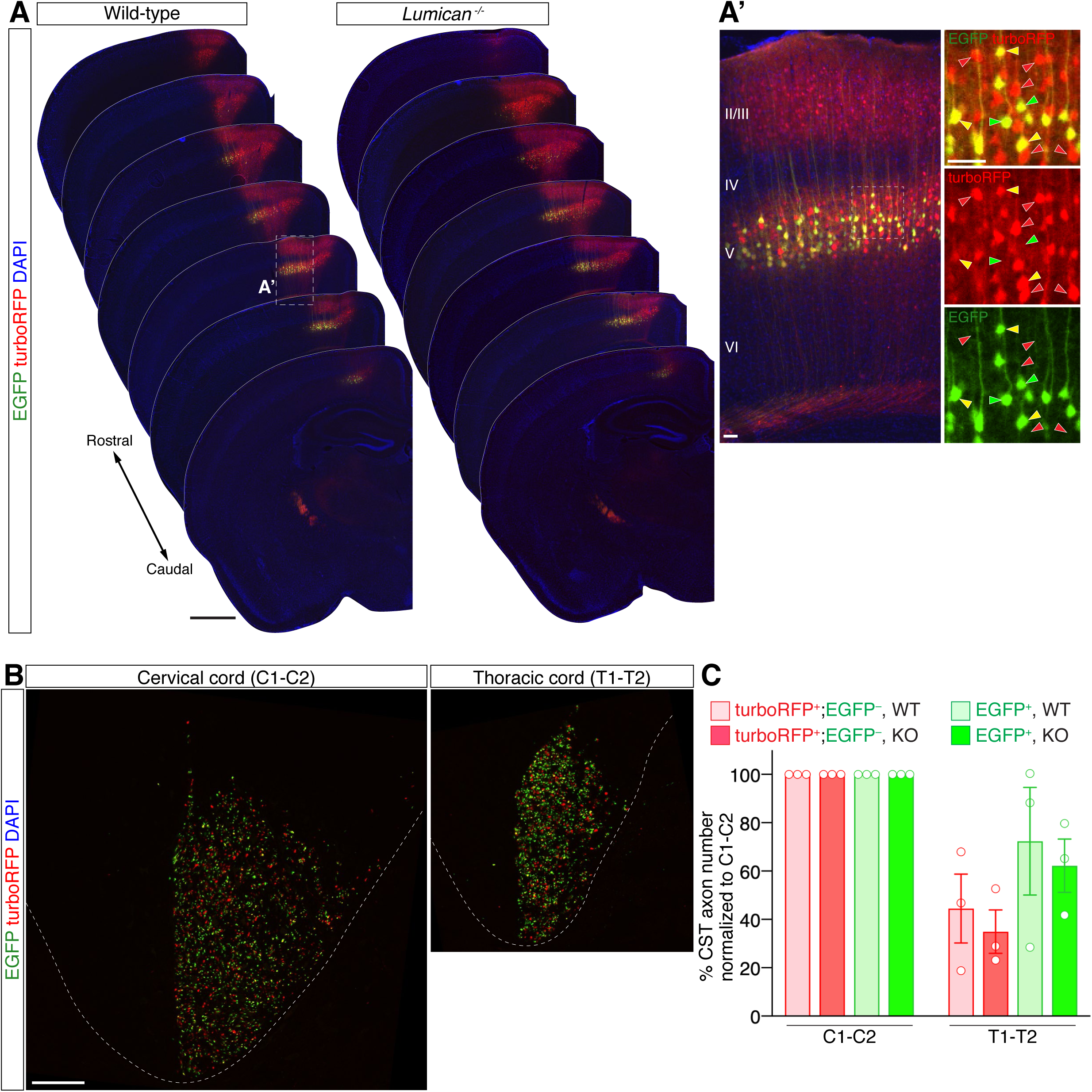
Exclusionary, subtractive viral labeling efficiently enriches labeling of the CSN_BC-med_ subpopulation. (A) Serial coronal brain sections of 6-week-old wild-type and *Lumican* null mice injected with AAVs as described in Fig. 5B show turboRFP (red) and EGFP (green) fluorescence. Cortical injection site is between 3^rd^ and 4^th^ sections. Note rostrocaudal gradient of turboRFP/EGFP fluorescence ratio in M1 layer V: turboRFP is high rostrally and EGFP is high caudally. (A’) Within M1, turboRFP^+^ cells are found across cortical layers, while EGFP^+^ cells are exclusively found in layer V. In layer V, turboRFP^+^;EGFP^−^ cells (red arrowheads), turboRFP^−^;EGFP^+^ (green arrowheads), and turboRFP^+^;EGFP^+^ cells (yellow arrowheads) exhibit highly interdigitated positioning. DAPI, blue. Scale bars, 1 mm (A); 50 µm (A’). (B) Single-plane confocal images of axial sections of cervical C1–C2 or thoracic T1–T2 show turboRFP (red) and EGFP (green) fluorescence of CST axons interdigitated in the dorsal funiculus. Each punctum was classified as a turboRFP^+^;EGFP^−^ or EGFP^+^ CST axon for quantification in (C). Scale bar, 50 µm. (C) CST axon extension by distinctly labeled CSN to the cervical and thoracic spinal levels in wild-type and *Lumican* null mice. Data are presented as mean ± SEM, n = 3.

**Supplementary Figure S5:**
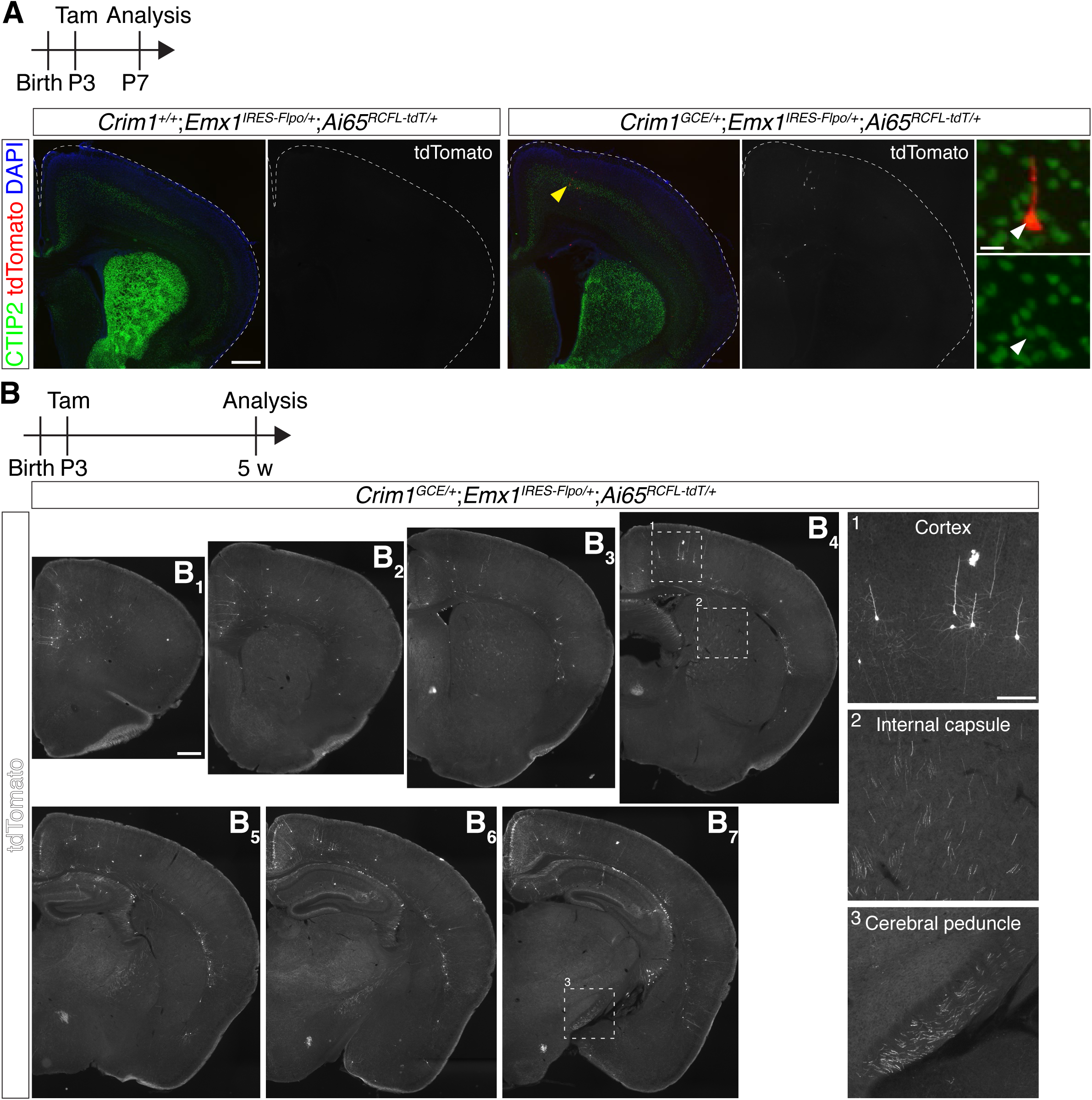
Intersectional genetic approach labels *Crim1*-expressing CSN. (A) Immunocytochemistry on coronal section of P7 *Crim1^+/+^;Emx1^IRES-Flpo/+^;Ai65^RCFL-tdT/+^* or *Crim1^GCE/+^;Emx1^IRES-Flpo/+^;Ai65^RCFL-tdT/+^* brains pulsed with tamoxifen at P3. While no tdTomato^+^ cells are found in *Crim1^+/+^* brains, *Crim1^GCE/+^* brains display rare labeled tdTomato^+^ cells (red). A tdTomato^+^ cell indicated by yellow arrowhead is magnified at right, showing co-localization with CTIP2 (green). Note that CTIP2-expressing tdTomato^+^ neurons are located medially, consistent with CSN_TL_ residing medially. DAPI, blue. Scale bars, 500 µm; 20 µm (inset). (B) Immunocytochemistry on serial coronal sections (B_1_–B_7_) of 5-week-old *Crim1^GCE/+^;Emx1^IRES-Flpo/+^;Ai65^RCFL-tdT/+^* brain pulsed with tamoxifen at P3 shows sparsely labeled tdTomato^+^ neurons in the cortex. Insets show tdTomato^+^ layer V neurons with thick basal dendrites in the cortex (1), and axon projections in the internal capsule (2) and cerebral peduncle (3), cardinal features of CSN. Scale bars, 500 µm; 200 µm (inset).

**Supplementary Figure S6:**
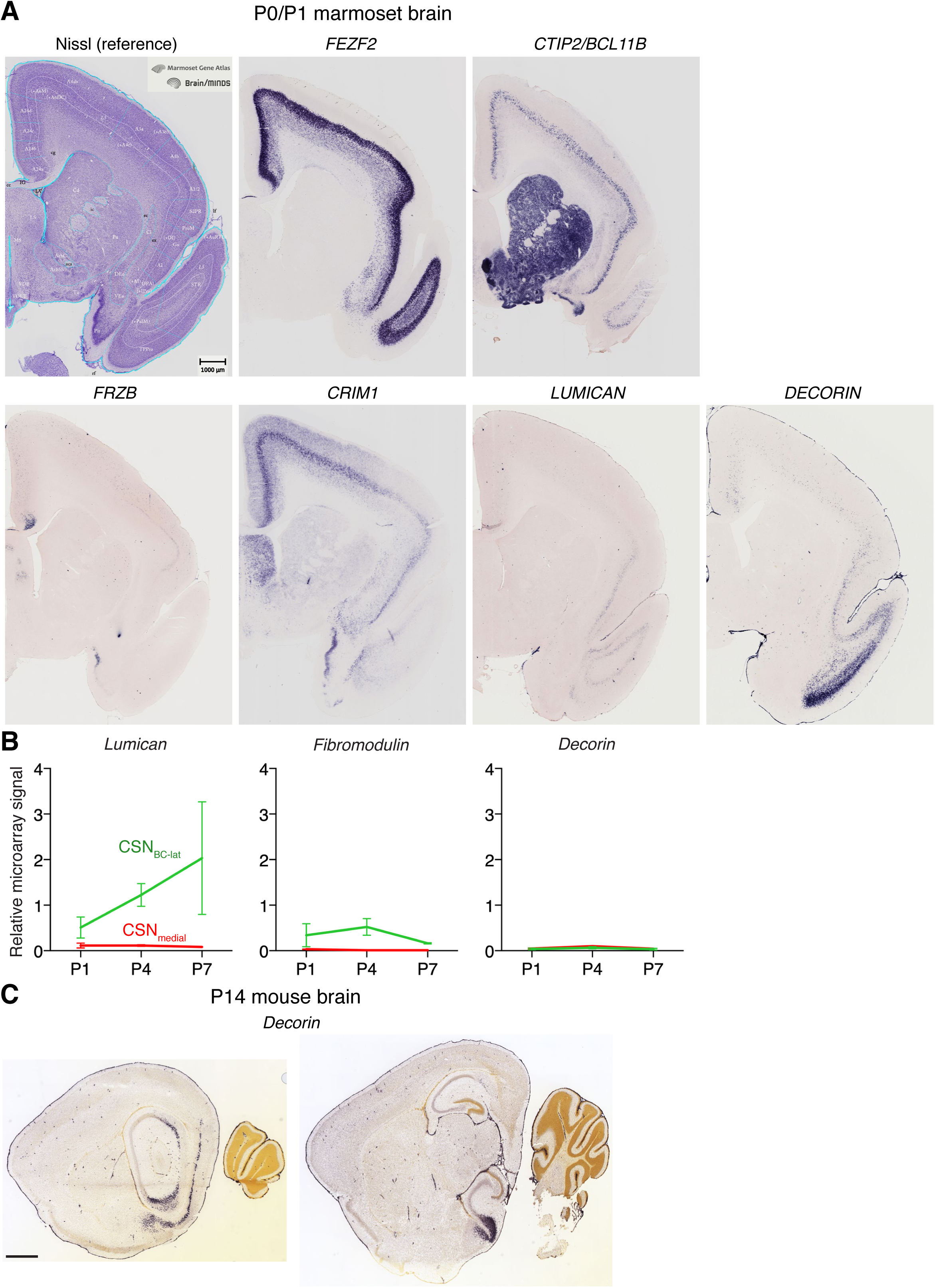
Expression of SLRP family genes in marmoset and mouse brains suggests the expanded use of SLRP family genes during mammalian CSN evolution. (A) Nissl staining and *in situ* hybridization for CSN/SCPN control genes *FEZF2* and *CTIP2*/*BCL11B*, CSN subpopulation-specific genes (*FRZB* for CSN_BC-lat_ and *CRIM1* for CSN_medial_/CSN_TL_, see (Sahni *et al*., 2018)), and SLRP family genes *LUMICAN* and *DECORIN* on P0/P1 marmoset coronal brain sections. *LUMICAN* is rostrolaterally expressed in layer V in marmoset cortex, consistent with our finding in mouse brain. Notably, *DECORIN* is also rostrolaterally expressed in layer V, with prominent ventral expression. P0/P1 marmoset brain developmentally corresponds to ∼P10 mouse brain (Clancy *et al*., 2001). All images shown here are taken from the Marmoset Gene Atlas website (https://gene-atlas.bminds.brain.riken.jp; Experiment #AI-1, #78-2, #140-4, #251-7, #262-5, #301-8) (Shimogori *et al*., 2018). Scale bar, 1 mm. (B) Temporal profile of SLRP family gene expressions from microarray analysis at postnatal ages P1, P4, and P7 in mice (Sahni *et al*., 2018). CSN_BC-lat_, green; CSN_medial_, red. *Lumican* panel is also shown in Fig. 1B. Besides *Lumican*, *Fibromodulin* is the only SLRP gene showing any significant differential expression: *Fibromodulin* expression peaks around P4, although at a much lower level than *Lumican*. *Decorin* does not show any significant, detectable expression. Y-axis represents normalized fluorescence intensity; data are presented as mean ± SD, n = 2–3. (C) *In situ* hybridization for *Decorin* on mouse P14 sagittal sections. *Decorin* expression is not observed in rostral cortex. Images are from the Allen Developing Mouse Brain Atlas website (Experiment: 100015428). Scale bar, 1 mm.

**Supplementary Figure S7:**
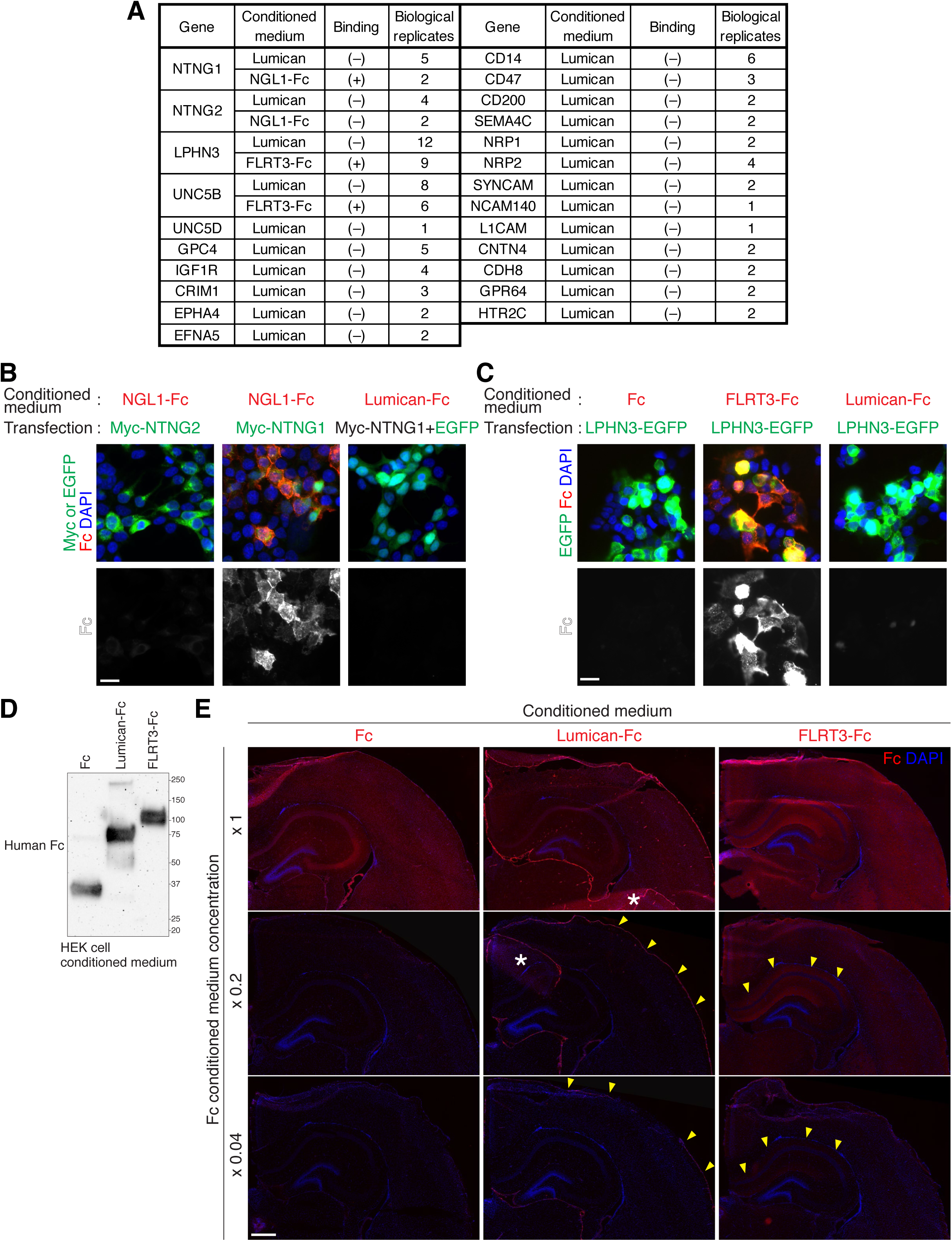
Cell surface and section binding assays reveal Lumican binding to meninges. (A–C) A HEK cell surface binding assay was used to assess potential Lumican interaction with select candidate molecules, listed in (A). HEK cells transiently transfected with one of the listed candidate molecules were incubated for 1 hour with the conditioned medium of HEK cells transiently expressing Lumican or Lumican-Fc fusion protein, and were subsequently immunolabeled with anti-Lumican or anti-Fc antibody to assess Lumican binding. Positive control ligand-receptor pairs (NGL1-NTNG1, LPHN3-FLRT3, and FLRT3-UNC5B) show substantial interaction between the ligand and receptor at the surface of transfected HEK cells, while a negative control ligand-receptor pair (NGL1-NTNG2) does not display any detectable interaction (B, C). None of the tested molecules exhibit any detectable interaction with Lumican above background (A–C). Scale bars, 20 µm. (D, E) A tissue section binding assay was performed to broadly assess potential Lumican interaction with cells or structures in the brain. The conditioned medium of HEK 293T cells transiently expressing either Fc, Lumican-Fc, or FLRT3-Fc (D) was applied to coronal sections of P10 *Lumican* null brain, followed by incubation for 1 hour and immunolabeling by anti-Fc antibody (E). While Fc does not show any binding at low concentrations, FLRT3-Fc displays binding to CA1 region in the hippocampus (arrowheads). Though Lumican-Fc binds efficiently to meninges (arrowheads), it does not exhibit interaction with any cortical cells at low concentrations. HEK 293T cells express Lumican core protein, but not the KSPG form. Asterisks indicate a fold in the section, with higher background. Scale bar, 500 µm.

**Supplementary Figure S8:**
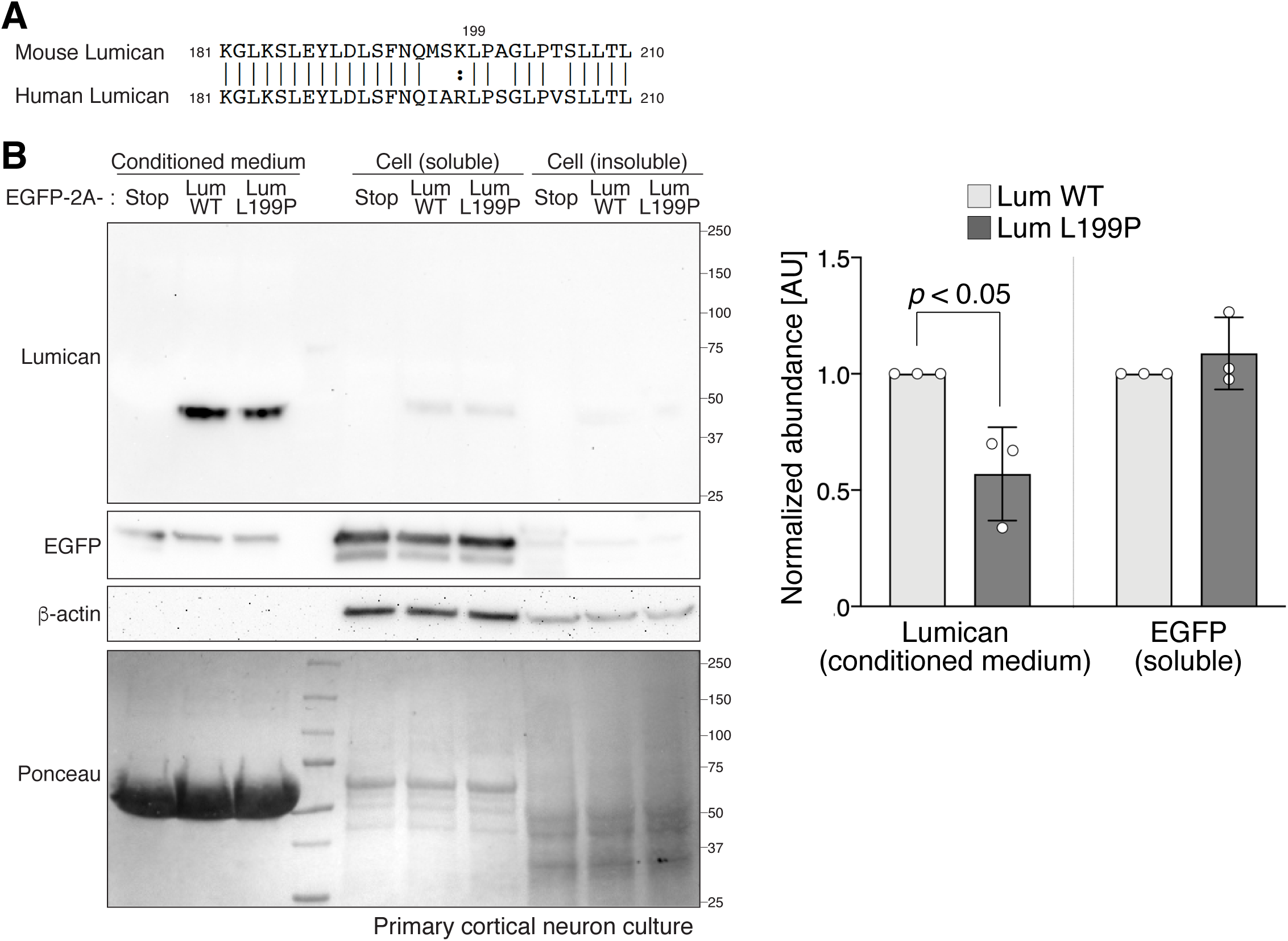
Mouse Lumican L199P mutant displays reduced protein abundance in the conditioned medium of primary cortical neuron culture. (A) Comparison of mouse and human Lumican protein sequences. Lumican L199 is conserved between mouse and human. (B) Immunoblot analysis of Lumican secretion in primary cortical neuron culture. Primary neurons derived from P0 mouse cortex were nucleofected with a plasmid encoding EGFP-2A-Stop, EGFP-2A-mouse wild-type Lumican (Lum WT), or EGFP-2A-mouse Lumican L199P mutant (Lum L199P), and cultured for 7 days *in vitro*, followed by collection of conditioned medium, soluble cellular fraction, and insoluble cellular fraction. 1.9%, 24%, and 24% of total conditioned medium, soluble cellular fraction, and insoluble cellular fraction was loaded, respectively. Both wild-type and L199P mutant Lumican are efficiently secreted. Notably, the abundance of Lumican L199P mutant is significantly lower than wild-type Lumican in the conditioned medium, while EGFP abundance in the cytosol is comparable between samples expressing wild-type and L199P mutant Lumican. Lumican protein is barely detectable in the insoluble cellular fraction. These results suggest that L199P variance in humans destabilizes Lumican protein, and does not cause its aggregation in cells. Primary cortical neurons express Lumican core protein, but not the KSPG form. Data are presented as means ± SD, n = 3.

